# RNArefine: AI-guided Atomic-Level Refinement of RNA Structures

**DOI:** 10.64898/2026.06.26.734804

**Authors:** Sho Tsukiyama, Yang Li, Kengo Sato, Hiroyuki Kurata, Yang Zhang

**Author notes:** All correspondence should be addressed to Yang Zhang.

## Abstract

Considerable progress has been made in AI-driven RNA structure prediction, but the resulting models often lack complete atomic details or suffer from severe stereochemical distortions and incorrect local interactions. We present RNArefine, an AI-guided hierarchical framework for atomic-level RNA structure refinement. RNArefine first predicts base-pairing and base-stacking interactions using geometric attention networks and then integrates the interactions with physics-based force fields to guide a two-step refinement strategy consisting of Monte Carlo conformational sampling followed by L-BFGS energy optimization. Large-scale benchmark experiments on both sequence-based prediction models and cryo-EM–derived structures demonstrated that RNArefine consistently improves stereochemical quality, interaction fidelity and physically penalized structural accuracy while preserving global topology. When applied to blind CASP16 RNA prediction models, RNArefine improved ranking scores for 28 of the top 30 groups. These results establish RNArefine as a robust open-source framework for transforming raw RNA folds into physically realistic atomic models for downstream structural and therapeutic applications.

## INTRODUCTION

RNA structures play central roles in cellular regulation, catalysis, and molecular recognition, while also serving as important therapeutic targets and modalities^1–5^. Accurate determination of RNA three-dimensional (3D) structures is therefore essential for understanding RNA function and enabling structure-guided therapeutic design. However, experimental RNA structure determination remains technically challenging and low-throughput, resulting in a substantial gap between known RNA sequences and experimentally resolved structures. This limitation has driven rapid advances in computational RNA 3D structure prediction.

Various computational approaches for RNA 3D structure prediction have been developed, ranging from template-based (e.g., ModeRNA^6^ and RNABuilde^7^) and knowledge-based methods (FARFAR2^8^, Vfold3^9^, and RNAComposer^10^) to recent deep learning frameworks such as DRfold^11^, DRfold2^12^, DeepFoldRNA^13^, trRosettaRNA^14^, RhoFold^15^, and AlphaFold3^16^. Although these methods have shown promising performance in predicting global RNA topology, many prediction pipelines still rely heavily on coarse-grained or backbone-centered structural representations and therefore either lack atomic-level structural details or contain substantial local geometric distortions, necessitating additional refinement to improve stereochemical accuracy and atomic-level interactions. Such refinement facilitates the generation of physically realistic RNA conformations by resolving steric clashes and accurately modeling nucleotide-specific interactions such as base-pairing and base-stacking^17^. This refinement is particularly important for downstream structural and functional applications, including docking simulations and virtual screening, where physically unrealistic models containing severe steric clashes can lead to unstable energy calculations or failure of subsequent simulations^18–20^. Physically realistic full-atom RNA models are therefore essential for RNA-targeted drug discovery, enabling reliable molecular simulations and robust evaluation of ligand binding and molecular recognition.

To address these limitations and improve the structural usability of predicted RNA models, several RNA refinement approaches have been developed. MD-based refinement is widely used because it enables structural relaxation under physics-based force fields^11, 15^, but unconstrained MD simulations may disrupt global topology, particularly for models containing severe stereochemical distortions and steric clashes. Additional RNA-specific refinement methods, including Arena^21^, QRNAS^22^ and RNAfitme^23^, improve local geometry and nucleotide conformations through geometry optimization and knowledge-based restraints. However, existing approaches still face substantial challenges in simultaneously preserving global topology, resolving stereochemical violations, and accurately modeling RNA-specific interactions. These limitations remain a major barrier to generating physically realistic and functionally reliable RNA structures for downstream structural biology and therapeutic applications.

In this study, we present RNArefine, an integrative framework that combines AI-predicted RNA interaction restraints with physics-based potentials and conformational sampling and optimization to address the long-standing challenge of atomic-level RNA structure refinement (**Fig. 1**). Rather than simply adjusting local atomic coordinates, RNArefine explicitly balances stereochemical correction, RNA-specific interaction formation, and preservation of the global fold. It first predicts base-pairing and base-stacking interactions directly from coarse-grained RNA geometries and then incorporates these interactions as probabilistic restraints during Monte Carlo sampling and L-BFGS energy minimization. This integration enables RNArefine to correct steric clashes and local geometric distortions while avoiding the topology distortions that often occur in conventional molecular dynamics–based refinement. Across multi-level benchmarking using experimentally determined RNA structures, Cryo-EM–derived models, and blind CASP16 predictions, RNArefine consistently improved stereochemical quality, interaction fidelity, and physically penalized structural accuracy at atomic resolution. Compared with existing refinement approaches, RNArefine achieved substantially greater reductions in steric clashes and stereochemical violations while maintaining global structural accuracy. Together, these results establish RNArefine as a general refinement framework for converting predicted RNA folds into physically realistic and functionally usable atomic models.

**Figure 1.**
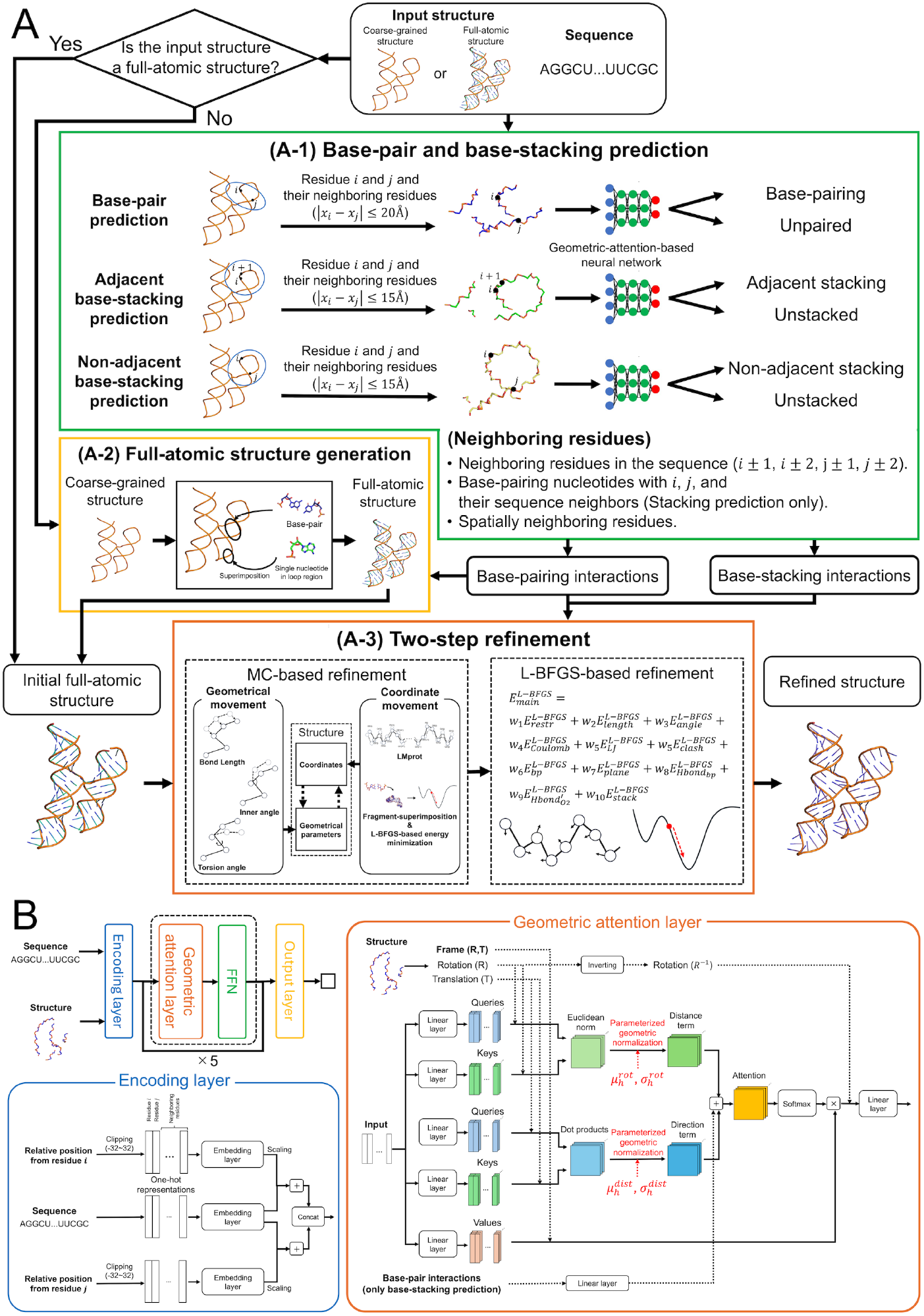
Workflow of RNArefine. (A) Overview of the RNArefine pipeline. (A-1) RNArefine predicts interactions using geometric attention–based neural network models. (A-2) For coarse-grained input models, full-atom RNA structures are reconstructed through fragment superimposition guided by the predicted interaction patterns. (A-3) Refinement is conducted via Monte Carlo simulations and L-BFGS energy minimization, using the predicted interactions as restraints. (B) Architecture of the geometric attention–based neural network model used for base-pair and base-stacking prediction.

## RESULTS

RNArefine consists of three major stages: interaction prediction, full-atom structure construction, and physics-guided refinement (**Fig. 1A**). Starting from coarse-grained RNA geometries, RNArefine first predicts base-pairing and base-stacking interactions directly from C4′-trace or backbone-frame representations using geometric attention–based neural networks (**Fig. 1B**). These predicted interactions are subsequently incorporated as probabilistic restraints during refinement. For coarse-grained input models, RNArefine reconstructs full-atom RNA structures through fragment superimposition guided by the predicted interaction patterns. The resulting full-atom models are then refined using a two-step strategy that combines Monte Carlo conformational sampling with L-BFGS energy minimization. By integrating AI-predicted interaction restraints with physics-based potentials, RNArefine simultaneously improves stereochemical realism, RNA-specific interaction fidelity, and preservation of the global fold.

### RNArefine accurately and robustly predicts base-pair and base-stacking interactions

#### Robust interaction prediction across native and predicted RNA structures

We first evaluate the predictive performance of RNArefine on base-pair and base-stacking interactions, which provide key restraints for downstream refinement. To evaluate different refinement scenarios, we assessed the ability of RNArefine to predict base-pair and base-stacking interactions in both native and predicted RNA 3D structures using our benchmark and Cryo-EM dataset (see **“Dataset construction”** in **Methods**). The predicted interactions were then compared with the corresponding ground truth.

The predicted structures were generated using two sequence-based methods (DRfold2 and AlphaFold3) for our benchmark dataset and a Cryo-EM density map–based method (EMRNA^24^) for a Cryo-EM dataset. AlphaFold3 and EMRNA directly produce full-atom RNA structures, whereas DRfold2 without post-processing generates coarse-grained models. These structures exhibited different levels of geometric quality, as assessed by clashscore and RMSD of bond lengths and bond angles. Compared with native structures, AlphaFold3 showed slightly distorted geometries, whereas EMRNA exhibited the largest geometric distortion (**Tables S1** and **S2** in **Supplementary Information, SI**). To characterize the performance of RNArefine, we compared it with the five existing methods (ClaRNA^25^, DSSR^26^, RNAView^27^, MC-Annotate^28^, and CSSR^29^) for base-pair identification and three methods (ClaRNA, DSSR and MC-Annotate) for base-stacking identification.

As shown in **Fig. 2**, and **Tables S3**-**S4**, in native structures of our dataset, the backbone frame-based prediction achieved F1-scores of 0.890 and 0.895 for base-pair and base-stacking prediction, respectively, whereas the C4′-trace–based prediction achieved 0.834 and 0.806. In the comparison of different base-pair types, canonical base pairs were predicted with higher F1-score and MCC values than non-canonical base pairs in both datasets, reflecting the diverse conformations of non-canonical base pairs arising from multiple combinations of interacting edges (Watson–Crick, Hoogsteen, and sugar edges) and relative orientations, as well as the limited number of training samples (**Tables S3** and **S5**). A similar trend was observed for base-stacking interactions, where the performance was higher for adjacent stacking than for non-adjacent stacking as non-adjacent base-stacking interactions allow a wider range of relative orientations than adjacent stacking (**Tables S4** and **S6**).

**Figure 2.**
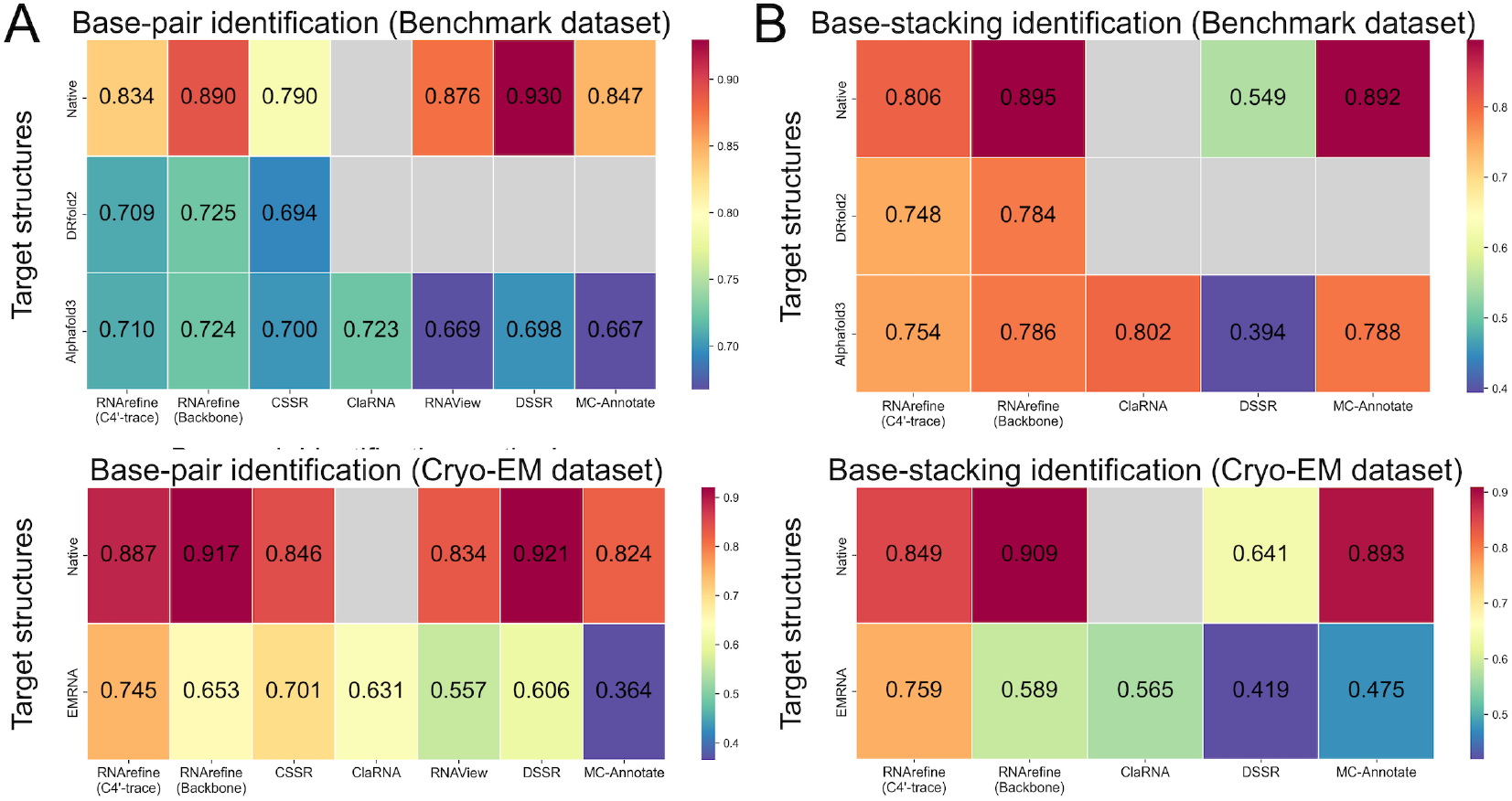
Performance comparison of base-pair and base-stacking identification on benchmark and Cryo-EM datasets. (A) Heatmaps of F1-scores for base-pair identification evaluated on our benchmark dataset (top) and the Cryo-EM dataset (bottom) across different types of structures and methods. (B) Corresponding heatmaps of F1-scores for base-stacking identification on our benchmark dataset (top) and the Cryo-EM dataset (bottom). Control methods include CSSR, ClaRNA, RNAView, DSSR, and MC-Annotate for base-pair prediction, and ClaRNA, DSSR, and MC-Annotate for base-stacking prediction. For both base-pair and base-stacking identification, the native (ground-truth) interactions were generated using ClaRNA applied to the native full-atom structures. Note that DRfold2 without refinement generates backbone-only structures; therefore, full-atom structure–based methods such as ClaRNA, RNAView, DSSR, and MC-Annotate are not applicable to DRfold2 predictions (empty gray cell).

For predicted structures, C4′-trace-based prediction demonstrated robust performance across different predicted structures. For example, although backbone frame-based prediction outperformed C4′-trace–based prediction for native structures, the two approaches showed comparable performance in structures predicted by AlphaFold3 and DRfold2 (**Fig. 2** and **Tables S7-10**). Furthermore, for structures predicted by EMRNA in the Cryo-EM dataset, C4′-trace-based prediction achieved F1-scores of 0.745 and 0.759 for base-pair and base-stacking prediction, respectively, greatly outperforming the backbone-frame–based prediction, which achieved F1-scores of 0.653 and 0.589, respectively (**Fig. 2** and **Tables S11** and **S12**). The superior performance of the C4′-trace-based approach in predicted structures likely arises because it does not require detailed atomic-level information and is therefore less sensitive to local geometric distortions.

#### Geometric attention improves interaction prediction robustness on distorted RNA structures

In the geometric attention-based neural network, we incorporated parameterized geometric normalization into the attention computation to enable each head to selectively respond to specific distance ranges and relative orientations and capture interaction-relevant geometric patterns. To evaluate its effect, we compared models with and without parameterized geometric normalization.

As shown in **Figs S1**-**S2** and **Tables S13**-**S14** in **SI**, parameterized geometric normalization generally improved performance when C4′-trace inputs were used, where the attention was computed by distance information only. In contrast, no significant improvement was observed with backbone-frame representations, which incorporate both distance and directional features. When only distance information is available, distance-based selectivity enhances interaction discrimination. On the other hand, when directional information is available, relative orientation may already provide sufficient information to distinguish interactions, reducing the additional benefit of explicit distance selectivity.

#### RNArefine outperforms existing interaction-identification methods on predicted structures

In base-pair prediction, DSSR achieved the highest performance on native structures, with F1-scores of 0.930 on our benchmark dataset and 0.921 on the Cryo-EM dataset (**Fig. 2** and **Tables S3** and **S5**). However, its performance was substantially reduced when applied to predicted structures, with F1-scores of 0.698 and 0.606 for AlphaFold3-predicted and EMRNA-predicted structures, respectively (**Tables S9** and **S11**). In contrast, our backbone-frame-based prediction demonstrated robust performance not only for native structures, achieving an F1-score of 0.890 on our benchmark dataset and a F1-score of 0.917 on the Cryo-EM dataset (**Tables S3** and **S5**), but also for predicted structures, where it showed high performance, with F1-scores of 0.725, 0.724, and 0.653 for DRfold2-, AlphaFold3-, and EMRNA-predicted structures, respectively (**Fig. 2** and **Tables S7, S9** and **S11**).

Furthermore, despite relying solely on backbone information, it outperformed full-atom structure-based methods such as RNAView, ClaRNA and MC-Annotate in both native and predicted structures (**Fig. 2** and **Tables S3, S5, S9**, and **S11**). The C4′-trace-based prediction showed the highest performance in the structure predicted by EMRNA (**Table S11**). In addition, it consistently achieved higher accuracy than CSSR, which is a P-trace-based method, across almost all prediction settings (**Tables S3, S5, S7, S9**, and **S11**). In base-stacking prediction, our method exhibited a similar trend to that observed for base-pair prediction (**Fig. 2** and **Tables S4, S6, S8, S10**, and **S12**). While ClaRNA and MC-Annotate achieved high performance on native structures, they showed limited ability in predicted structures.

Although our models were trained on native structures, they achieved high performance in interaction identification even for predicted structures. This robustness likely arises from the combination of the coarse-grained structural representation and the geometric attention-based neural network. Unlike existing methods that emphasize interatomic distances and detailed base-region geometry, our approach focuses on residue-level spatial and sequential relationships, enabling robust predictions even under distorted or degraded geometries. The geometric attention mechanism further captures RNA-specific topological interaction patterns, contributing to higher performance on both native and predicted structures. These results demonstrate its applicability to base-pair and base-stacking prediction prior to structural refinement.

### RNArefine refines RNA structures toward native-like atomic quality

For the evaluation of refinement with our benchmark dataset, DRfold2 was used for generating initial structures. Since DRfold2 generates coarse-grained structures (backbone atoms + N9/N1), full-atom structures were constructed by superimposing two types of fragments, base-pair fragments and single nucleotide fragments in loop regions, onto the structures (see **“Full-atom structure generation”** in **Methods**). The selection of fragment types was guided by the predicted base-pair information. To assess our refinement comprehensively, 16 metrics including RMSD, TM-score, penalized TM-score, Clashscore, lDDT, lDDT with stereochemical quality checks (hereafter Penalized lDDT), RMSD (bond length), RMSD (bond angle), MolProbity score, HB-score, MCQ, INF_all, INF_stack, INF_wc, INF_nwc, and GDT-TS were calculated (see **“Evaluation metrics for structure refinement”** in **Methods**). Here, to rigorously evaluate structural accuracy and prevent models from achieving high topological scores through unphysical atomic geometries, we employed a penalized TM-score as defined in Eq. (35). This stereochemistry-aware evaluation was originally introduced in the lDDT metric^30^ and later adopted for RNA structure evaluation by CASP assessors^31^.

As shown in **Fig. 3A** and **Table S15**, compared with the initial full-atom structures, RNArefine improved 13 out of 16 metrics. Importantly, TM-score, GDT-TS and RMSD values remained largely unchanged after refinement, indicating that the global folding topology was well preserved. In contrast, substantial improvements were observed in geometric quality. For example, the clashscore was dramatically reduced from 116 in the initial full-atom structures to 8.16 after refinement. Similarly, RMSD from ideal bond lengths and bond angles decreased from 0.0470 Å and 8.28° to 0.0143 Å and 1.67°, respectively. These reductions demonstrate a marked improvement in physical realism.

**Figure 3.**
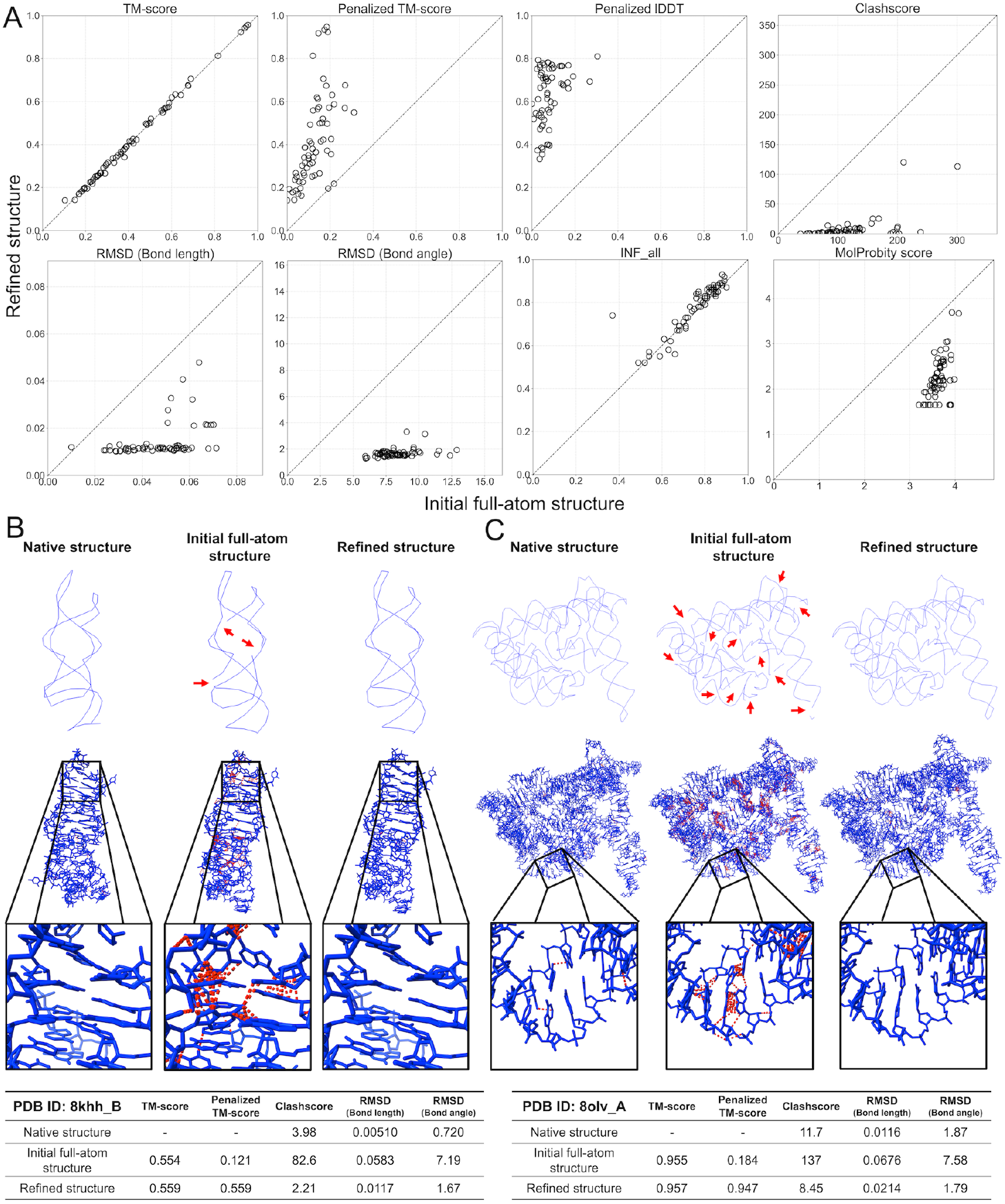
Comparison between structures before and after refinement using RNArefine. (A) Comparison of metrics between the initial full-atom structures and the refined structures. Scatter plots show TM-score, penalized TM-score, lDDT, Clashscore, RMSD of bond lengths, RMSD of bond angles, INF, and MolProbity score. Each point corresponds to one target RNA, and dashed lines indicate equal performance before and after refinement. (B) Representative example of structure refinement for the 2’-dG-III riboswitch with guanosine (PDB ID: 8khh, chain B). The native structure, the initial full-atom structure, and the refined structure are shown. The first row shows the backbone-traces, and the second row the full-atomic structures. Evaluation metrics, including TM-score, lDDT, Clashscore, and RMSDs of bond lengths and bond angles, are summarized below each example. Red lines within the structures in the second row indicate atomic clashes, where red arrows indicate regions with disconnected bonds in the initial backbone-trace structure. (C) Same as (B), but for the Oceanobacillus iheyensis group II intron (PDB ID: 8olv, chain A).

Structural inspection further confirms these improvements, showing that RNArefine generates covalently continuous structures while effectively resolving steric clashes between neighboring residues (**Figs. 3B-C**). The clashscore after refinement was comparable to that of the native structures, indicating that the refined models achieve near-native stereochemical quality. Notably, the average MolProbity scores across samples for the RNArefine-refined structures were slightly better than those of the native structures (**Table S16**). Consistently, penalized TM-score and penalized lDDT, which incorporate penalties for geometric violations into TM-score and lDDT, respectively, showed substantial improvement after refinement. These results further support the suggestion that RNArefine enhances local geometric correctness without compromising global structural integrity.

In addition, RNArefine showed high scores in interaction network fidelity (INF). Notably, stacking interactions and non-Watson–Crick base pairs (INF_stack and INF_nwc) were substantially improved, likely due to the incorporation of interaction-specific energy terms during refinement. In contrast, INF_wc, which evaluates Watson–Crick base pairs, remained nearly identical to that of the initial full-atom models, as these models were constructed by superimposing base-pair fragments at positions predicted to form base pairs and therefore already formed Watson–Crick base pairs. Furthermore, the improvement in HB-score indicates enhanced hydrogen-bond formation, suggesting that RNArefine captures atomic-level physical constraints underlying RNA structure formation.

### Contribution of predicted interaction restraints and multi-step refinement

#### Predicted interaction restraints improve topology preservation and interaction fidelity

A critical feature of RNArefine is the use of interaction information predicted by geometric attention-based neural networks as probabilistic constraints during refinement. To evaluate the contribution of these predicted interactions to refinement, we compared the performance of the standard RNArefine pipeline with that of RNArefine without energy terms associated with RNA-specific interactions.

As shown in **Fig. S3** and **Table S17**, refinement without interaction information resulted in a lower clashscore than the full RNArefine pipeline. This is because removing interaction restraints may allow steric energy terms, such as those modeled by the Lennard-Jones potential, to play a more dominant role, thereby reducing steric clashes. On the other hand, refinement without interaction restraints reduced global structural metrics, including TM-score^32, 33^ and GDT-TS^34^, compared with the standard RNArefine pipeline.

In addition, the interaction network fidelity (INF), which measures the consistency of RNA-specific interactions with the native structure, was also significantly higher when interaction restraints were applied (sample-wise one-sided t-test). The HB-score was also improved when interaction information was incorporated, indicating that the formation of RNA interactions was enhanced even at the atomic level during refinement. These results suggest that predicted interaction restraints play an important role in improving RNA interaction fidelity and preserving global topology, thereby enabling more comprehensive refinement.

#### Monte Carlo sampling and L-BFGS minimization play complementary roles during refinement

In addition to the contribution of interaction restraints, we next examined the role of the multi-step refinement procedure in improving structural quality. RNArefine combines two complementary refinement strategies: Monte Carlo (MC) simulation and L-BFGS-based energy minimization. To evaluate the contribution of each step, the performance of the initial full-atom structures, the fully refined structures (RNArefine), and the structures refined by each step individually was compared. To jointly evaluate both global and local aspects of RNA structural quality using a single measure, we introduced a composite score that integrates standardized evaluation metrics (Z-scores) in four structural quality categories (see **Eqs 36–40**). Here, the composite score was calculated across the compared structure sets.

As shown in **Table S15**, MC-only refinement achieved a composite score of 0.808, and L-BFGS-only refinement achieved 1.19, whereas the combined two-step refinement achieved the best performance, with a composite score of 1.31. Consistent with the Z-score improvement, the combined refinement also showed significantly better geometric quality than the other conditions in terms of clashscore, RMSD (bond length), RMSD (bond angle), and MolProbity score, as assessed by sample-wise one-sided t-tests (**Fig. S4**).

These results indicate that MC simulation and L-BFGS minimization play complementary roles in refinement. Specifically, MC simulation is responsible for structural exploration and relaxation, while L-BFGS efficiently performs optimization along the energy landscape, resulting in more effective refinement than either method alone. Their combination therefore enables simultaneous optimization of stereochemical quality and global structural preservation.

### RNArefine achieves superior stereochemical refinement while preserving global topology

#### RNArefine achieves superior refinement performance across multiple structural quality metrics

The performance of RNArefine was compared with that of existing methods including PDBfixer, RNAfitme, Arena, QRNAS, and MD simulation based on the AMBER force field implemented in OpenMM 7.7, following the setup used in RhoFold++ (see **“Setting of existing refinement methods”** in **Methods** for detailed settings). To comprehensively evaluate refinement performance, we calculated a composite score for all methods. As shown in **Fig. 4** and **Table S18**, RNArefine achieved the best composite score and the highest performance in 11 of 16 evaluation metrics. In particular, RNArefine substantially improved geometric quality, achieving lower clashscores and smaller RMSDs from ideal bond lengths and angles than existing approaches across almost all samples (**Fig. 4B**). It also obtained the highest penalized lDDT and penalized TM-score values in most cases.

**Figure 4.**
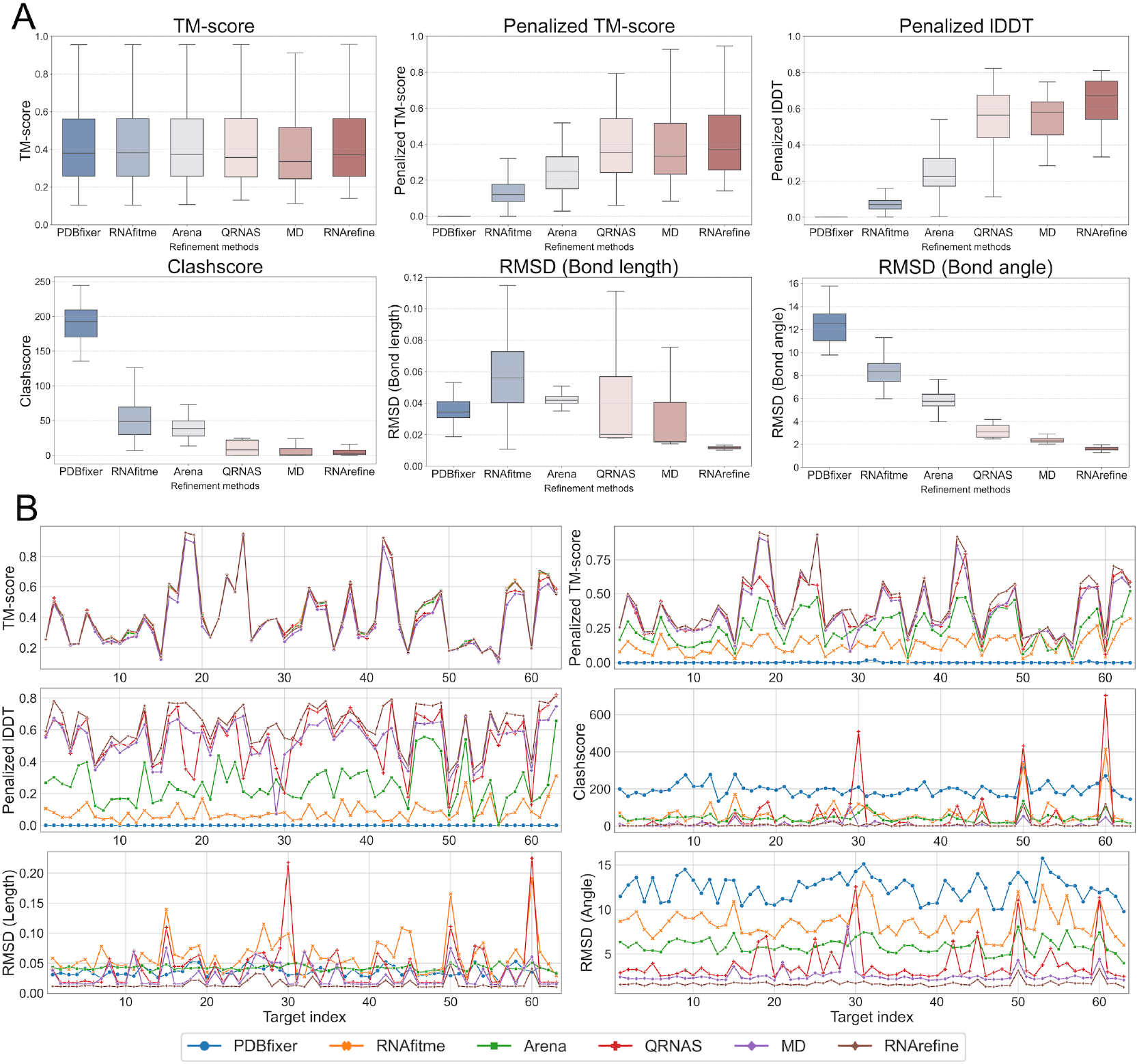
Benchmark evaluation of RNA structure refinement performance. (A) Box plots comparing refinement performance across different methods on a benchmark dataset. TM-score, penalized TM-score, lDDT, Clashscore, RMSD of bond lengths, and RMSD of bond angles are shown for each refinement method. Boxes indicate the interquartile range (IQR; 25th–75th percentiles) with the median shown as a center line; whiskers extend to the most extreme values within 1.5×IQR. (B) Per-sample refinement performance on the benchmark dataset. Each plot shows the corresponding metric for individual RNA targets.

#### Interaction-guided refinement and hybrid optimization improve topology preservation during refinement

Although Arena and RNAfitme enable rapid refinement, their refinement effects were limited. MD simulation achieved clashscores comparable to those of RNArefine. However, its performance in global folding metrics including TM-score, RMSD, and GDT-TS was inferior to those of the initial full-atom structure and RNArefine. For example, MD frequently reduced TM-scores across many targets, indicating disruption of global structural topology during refinement (**Fig. 4B**). In addition, MD resulted in poorer INF values, indicating reduced accuracy in intramolecular interaction formation. These results suggest that while MD improves local geometric quality, it may disrupt base-pair and base-stacking interactions and distort global folding during refinement. These limitations substantially reduced the overall refinement performance of MD (**Table S18**). QRNAS addresses this limitation of MD by introducing restraint potentials that prevent the distortion of the global fold during geometric refinement, resulting in the second-highest composite score following RNArefine.

In RNArefine, we adopted a similar restraint strategy and further incorporate fragment-based energy terms guided by predicted base-pair and base-stacking interactions. Moreover, whereas QRNAS relies solely on steepest-descent energy minimization, RNArefine employs L-BFGS optimization combined with Monte Carlo sampling to explore diverse local conformational space. Through this combination of knowledge-based restraints and advanced conformational sampling, RNArefine avoids the structural distortions observed in MD-based refinement and improved physical realism, resulting in a markedly higher composite score compared with existing approaches.

### RNArefine restores physical realism in Cryo-EM–derived RNA structures

In recent years, deep learning–based approaches have been developed to identify RNA structures directly from cryo-EM density maps. While these methods are effective at capturing backbone structure, the resulting models often lack sufficient physical realism and accurate intramolecular interactions. To address this limitation, we applied RNArefine to the RNA structures predicted by EMRNA^24^, a deep learning–based method for structure prediction from cryo-EM density maps. The predicted models were collected from the main test dataset in the original EMRNA study, and after removing structures overlapping with our training dataset, 15 targets were selected for refinement.

#### RNArefine substantially improves physical realism and topology of Cryo-EM–derived RNA models

The Cryo-EM-predicted structures contain substantial physical violations (**Table S2**). Therefore, before refinement, base-pairing and base-stacking interactions were predicted not only from backbone frames but also from the C4′ trace. As shown in **Fig. 5** and **Table S19**, in almost all metrics, both backbone-based RNArefine and C4′-trace-based RNArefine led to consistent improvements over the original EMRNA models. Notably, although these refinements were performed without directly using cryo-EM density maps, RNArefine improved not only local stereochemical metrics, such as clashscore, but also global topology, such as TM-score. The improvement in global topology may result from the correct assembly of base-pair and base-stacking interactions. Because INF scores increased across all interaction types in the refined structures (**Table S19**), base-pair and base-stacking interactions were more accurately formed. This interaction formation contributed to improvements in global topology.

**Figure 5.**
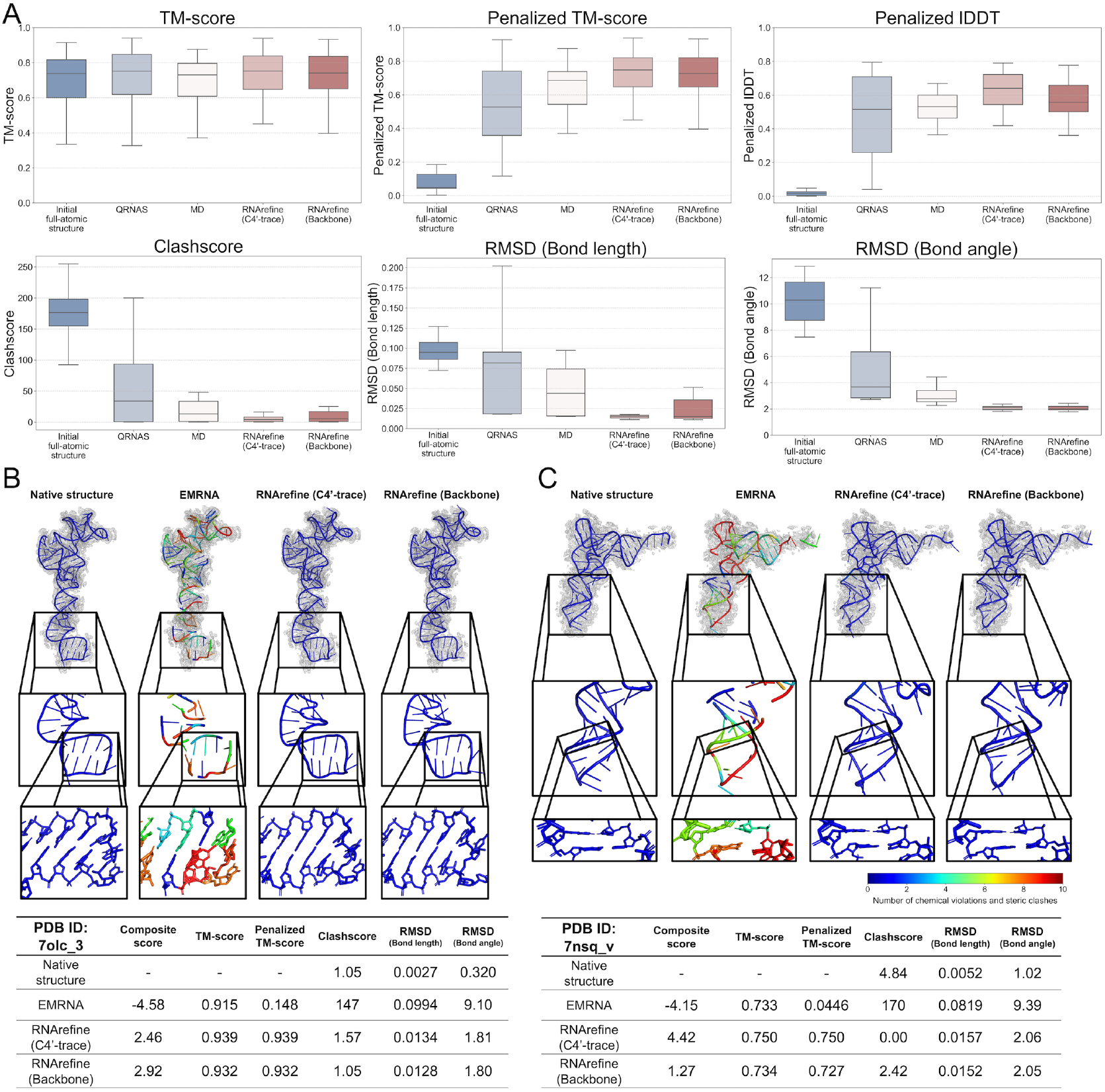
Evaluation of refinement using Cryo-EM–determined structures. (A) Box plots comparing metrics for different refinement methods evaluated using Cryo-EM–determined structures. TM-score, penalized TM-score, lDDT, and Clashscore are shown for structure-based refinement methods (RNArefine, QRNAS, and MD). Boxes indicate the interquartile range (IQR; 25th–75th percentiles) with the median shown as a center line; whiskers extend to the most extreme values within 1.5×IQR. (B) Representative refinement example for the 5S rRNA in the Thermophilic eukaryotic 80S ribosome (PDB ID: 7olc, chain 3). The native structure, the initial full-atom structure, and refined structures are shown. The structures are colored based on the total number of steric clashes and chemical violations (including bond lengths and bond angles) identified by the OpenStructure toolkit^47^. The color scale is capped at 10, where values of 10 or higher are shown in maximum red. Evaluation metrics are summarized in the tables. (C) Similar to (B) but for the AtRNA (Arg) in the E. coli ribosomal complex with A and P-tRNA (PDB ID: 7nsq, chain v).

Moreover, the structures were refined with respect to stereochemical quality and physical realism. For example, the clashscore dropped dramatically from 176 in the original EMRNA models to 9.04 and 5.53 after refinement with backbone-based RNArefine and C4′-trace-based RNArefine, respectively. In addition, bond length and bond angle deviations (RMSD from ideal bond lengths and bond angles) were also substantially reduced. As shown in **Figs. 5B** and **5C**, EMRNA models showed fragmented structures with broken covalent connections, whereas the refined models exhibited continuous covalent chains. Consistent with the visual inspection, the refined structures exhibited near-native physical quality, as assessed by Clashscore and MolProbity score (**Figs. 5B** and **C**, and **Table S20**).

#### C4′-trace–guided refinement is particularly robust for geometrically distorted RNA structures

In the refinement of EMRNA-predicted structures, which often exhibit geometric inconsistencies and stereochemical violations, C4′-trace-based RNArefine achieved superior performance across multiple metrics compared with the backbone-frame-based approach, suggesting more reliable prediction of base-pairing and stacking interactions from C4′-trace information (see **“RNArefine accurately and robustly predicts base-pair and base-stacking interactions”**). These findings suggest that the accuracy of interaction prediction directly influences downstream refinement quality, and that C4′-trace–guided refinement is particularly robust when starting from geometrically distorted initial models.

#### RNArefine outperforms MD- and restraint-based refinement methods on Cryo-EM structures

We additionally refined the EMRNA-generated models using MD simulations and QRNAS, and composite scores were calculated to compare RNArefine with these methods. Both backbone-based RNArefine and C4′-trace-based RNArefine substantially outperformed QRNAS (composite score = 0.240) and MD simulation (composite score = −0.805), achieving composite scores of 3.26 and 1.58, respectively. Consistent with the observations from our benchmark evaluation (see **“RNArefine achieves superior stereochemical refinement while preserving global topology”**), MD-based refinement improved certain geometric metrics, for example by reducing the clashscore from 176 to 28.2, but disrupted global topology, resulting in lower TM-score and GDT-TS values and higher RMSD.

QRNAS enhanced INF values and facilitated the formation of base-pair and base-stacking interactions, leading to improved global topology. However, its performance in local stereochemical quality remained inferior to RNArefine. RNArefine not only refined the original EMRNA models but also outperformed both MD and QRNAS in nearly all evaluation metrics, improving both local stereochemistry and global structural accuracy. These results suggest that RNArefine overcomes limitations of existing refinement methods and has the potential to facilitate automated determination of RNA 3D structures with physical realism and structural fidelity from experimental data.

### RNArefine consistently improves blind CASP16 RNA prediction models

#### RNArefine improves prediction scores across CASP16 participant models

To evaluate the performance of RNArefine on diverse initial structures, we applied it to the models submitted to CASP16 by multiple groups. Sixteen RNA monomer targets whose experimental structures are available in the PDB were used for this evaluation (see **“Dataset construction” in Methods**). Refinement was applied to the first models of the top 30 groups on these 16 targets. Before refinement, base-pair and base-stacking interactions were predicted from the backbone frames of these models, and the structures were subsequently refined.

According to the standard CASP16 ranking policy for RNA monomers, the overall score is evaluated by combining three metrics: TM-score, GDT-TS, and lDDT, with weights of 0.3, 0.3, and 0.4, respectively. Since official CASP native structures can sometimes differ from standard PDB entries, we estimated the CASP-equivalent scores by adjusting the official CASP scores based on score differences computed against the PDB structures (see **Eq. 41**). Following the standard CASP16 ranking policy^31^, we computed Z-scores for each metric across the top 30 groups while excluding outliers. The final combined Z-score for each target was calculated as 0.3 × *Z*_*TM-score*_ + 0.3 × *Z*_*GDT-TS*_ + 0.4 × *Z*_*lDDt*_, with any negative values set to zero.

As shown in **Figs. 6** and **S5** and **Tables S21** and **S22**, Z-scores improved for 28 of the top 30 groups after refinement. Notably, when focusing on the top 15 groups, performance improvements were consistently observed across all groups. Two groups (CoDock and RNADojo) did not show improvement in the final score after refinement.

**Figure 6.**
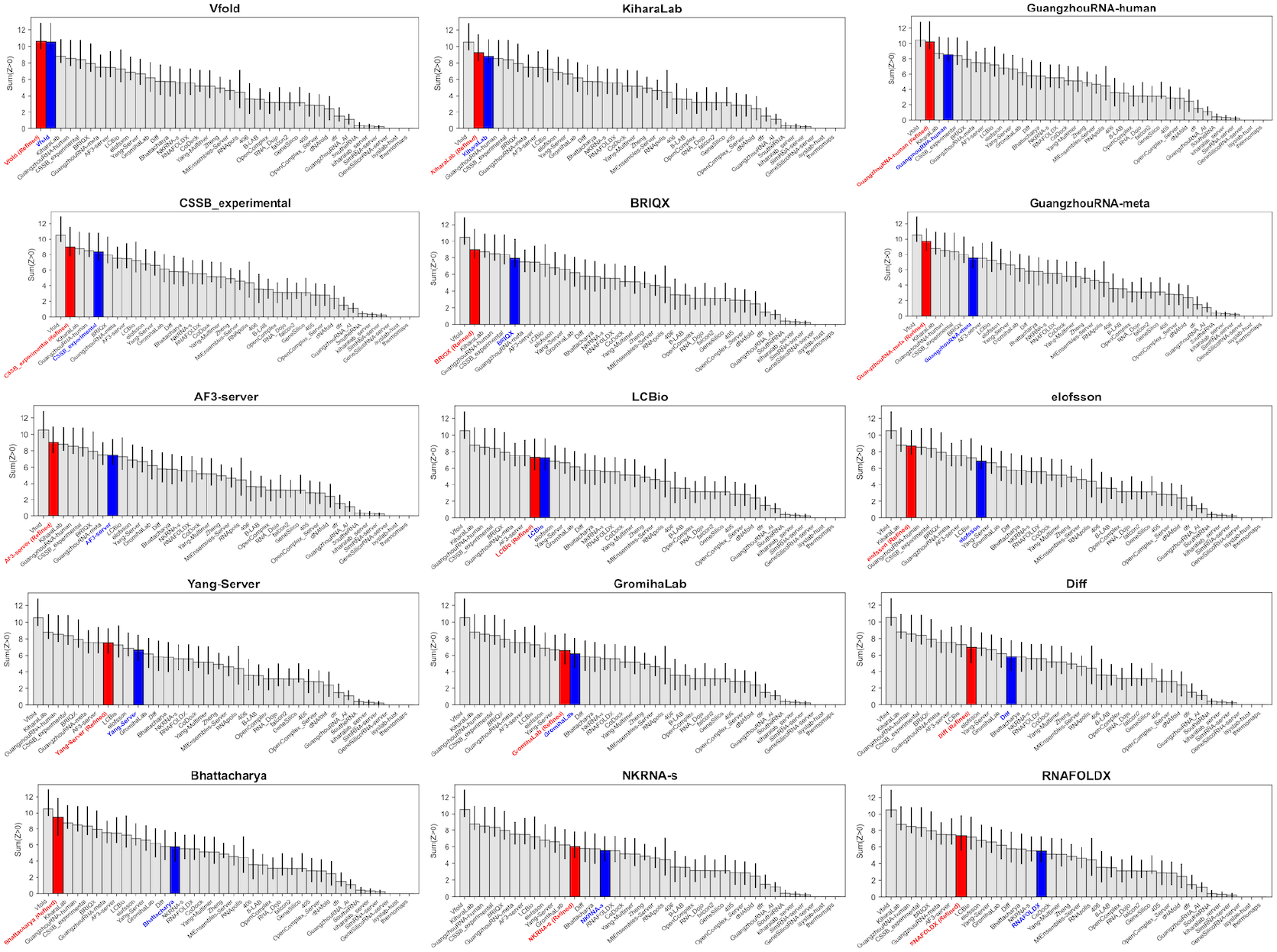
Improvement in RNA structure prediction performance by refinement in CASP16 (top 15 participant groups across 16 targets). Changes in CASP rankings before and after refinement are shown for the top 15 participating groups. Each panel corresponds to a participant group and displays the summed Z-score ranking across groups based on the originally submitted models (blue) and the estimated ranking after refinement (red). Z-scores and rankings were computed following the CASP assessment procedure, with confidence intervals estimated by bootstrap resampling using code adapted from the publicly available CASP16 nucleic acid assessment scripts^45^.

#### RNArefine complements accurate global folding by improving local structural quality

Importantly, the effect of refinement was observed for groups that achieved high scores in global fold metrics such as TM-score and GDT-TS but showed lower performance in local metrics and geometric quality, such as penalized lDDT.

For example, the group “Bhattacharya” ranked 7th in terms of the TM-score Z-score but dropped to 29th in the lDDT Z-score, resulting in an overall rank of 13th (**Table S22**). Similarly, while “dNAfold” achieved 19th in the TM-score Z-score, it ranked last (30th) in lDDT, leading to an overall rank of 29th. By applying RNArefine to the models from these groups, their rankings improved substantially, with Bhattacharya rising to 2nd place and dNAfold to 16th. Similar trends were observed across many groups showing gaps between global and local structural performance. In particular, the highest scores were achieved in the refined “Vfold” structures (**Table S21**).

CASP evaluates models comprehensively, considering not only global folding but also local structure and stereochemical correctness. Therefore, these results demonstrate that RNArefine effectively complements methods that predict accurate global folds yet exhibit limitations in local structure and geometric quality. In addition, these findings suggest that RNArefine serves as an effective post-prediction refinement strategy and substantially expands the potential of existing RNA 3D structure prediction methods.

## DISCUSSION

Accurate RNA 3D structure prediction has advanced rapidly with the emergence of deep learning–based predictors, but predicted models often remain limited by local stereochemical distortions and incorrect RNA-specific interaction patterns. To overcome these limitations of RNA 3D structure prediction, we developed RNArefine, a refinement framework that integrates interaction prediction from coarse-grained representations with a two-step refinement strategy combining MC sampling and L-BFGS energy minimization.

### Robust interaction-guided refinement from coarse-grained RNA representations

RNArefine demonstrated high performance in base-pair and base-stacking identification not only on native structures but also on predicted models. While some state-of-the-art RNA 3D structure prediction methods successfully predict global topologies, the predicted structures often contain substantial local distortions in base regions. Existing base-pair and base-stacking identification tools, which were built on high-resolution native structures, rely heavily on precise atomic geometries and therefore fail to accurately detect interactions in geometrically distorted predicted models. In contrast, RNArefine employs a geometric attention–based neural network with parameterized geometric normalization to capture interaction-relevant geometric patterns from coarse-grained structural representations, enabling robust identification of base-pair and base-stacking interactions. This robustness ensures that reliable interaction restraints are provided for the subsequent refinement step, even for structures with substantial geometric errors.

### Hybrid refinement improves stereochemical realism while preserving global topology

In the refinement stage, RNArefine achieved dramatic improvements in stereochemical quality while avoiding the structural degradation often observed in conventional MD-based refinement. In our benchmark evaluation, MD simulation demonstrated strong local refinement capability relative to other existing approaches. However, it frequently disrupted global topology, potentially because severe steric clashes and chemical violations generated large repulsive forces during unconstrained relaxation. In RNArefine, to suppress such effects and stabilize refinement, we introduced fragment-based energy terms that use predicted base-pair and base-stacking interactions as probabilistic restraints to promote their formation, together with an energy restraint that preserves the global fold of the initial structure.

Furthermore, the two-step strategy combining MC simulation and L-BFGS energy minimization enables extensive conformational exploration and efficient energy optimization. During MC refinement, conformations were explored not only in Cartesian coordinate space but also in geometric parameter space, including bond lengths, bond angles, and torsion angles. Such cross-space conformational exploration facilitates correction of complex stereochemical violations. Combined with L-BFGS energy minimization, this strategy enables efficient energy reduction while maintaining stable local convergence. Consequently, RNArefine achieved substantial improvements in physical realism, including reduction of the clashscore from 116 to 8.16 in our benchmark evaluation (**Fig. 3** and **Table S15**).

### Broad applicability across prediction pipelines and structural states

RNArefine improved not only DRfold2-predicted structures in our benchmark evaluation but also enhanced the scores of models submitted by 28 of the top 30 groups in CASP16. The consistent improvement suggests that RNArefine corrects geometric errors and local stereochemical violations that remain unresolved in current RNA structure prediction pipelines. Notably, the refinement effect was particularly evident for methods that achieved accurate global folds but showed limitations in local structural quality and stereochemical correctness.

In addition to sequence-based prediction pipelines, RNArefine successfully refined structures constructed directly from cryo-EM density maps. RNArefine substantially improved both local stereochemistry and global topology without directly using density information during refinement, indicating that interaction-guided refinement can effectively recover physically realistic RNA conformations even from highly distorted initial models.

Together, these results establish RNArefine as a general framework for transforming predicted RNA folds into physically realistic atomic models while preserving global topology. By integrating AI-predicted interaction restraints with physics-based refinement and conformational optimization, RNArefine addresses a long-standing challenge in atomic-level RNA structure refinement that has remained difficult for both protein and RNA modeling communities^35–37^. The robust performance across benchmark datasets, cryo-EM–derived structures, and blind CASP16 predictions further suggests broad applicability to modern RNA structure prediction pipelines. These advances may facilitate downstream applications in structural biology, molecular simulation, and RNA-targeted therapeutic design.

## METHODS

As illustrated in **Fig. 1A**, the RNArefine pipeline consists of three steps, including base-pair and base-stacking prediction, full-atom structure construction, and refinement. In preparation for the pipeline execution, geometric attention-based neural network models were developed for the prediction of base-pair and base-stacking interactions, and structure fragments and statistical parameters were derived from high-resolution experimental structures. During the RNArefine pipeline, base-pair and base-stacking interactions are predicted directly from the input structure and sequence using the geometric attention-based neural network (**Fig. 1**). When the input structure is a coarse-grained model, an initial full-atom structure is constructed by superimposing the structure fragments on the coarse-grained structure. The initial full-atom structure is then refined using MC simulation followed by L-BFGS-based energy minimization, guided by the hybrid potential function containing deep learning-predicted base-pair and base-stacking interaction restraints and physical potentials.

### Dataset construction

The training dataset of DRfold^11^ was used for construction of base-pair and base-stacking interaction prediction models, as well as for generating structure fragments and statistical parameters. This dataset includes 3864 RNA structures which were deposited in the Protein Data Bank (PDB)^38^ before 2021.

To annotate base-pair and base-stacking interactions in the tertiary structures, ClaRNA^25^ was used. Nucleotide pairs that were not identified as base-pairs but had a C4′–C4′ distance of ≤ 20 Å were labeled as non-base-pairs. Similarly, nucleotide pairs that were not identified as stacking interactions but had a C4′–C4′ distance of ≤ 15 Å were labeled as non-stacking pairs. Nucleotide pairs (*i, j*) with a sequence separation of |*i* − *j*| ≤ 2 were excluded from the non–base-pair category, since adjacent nucleotides are structurally unlikely to form base-pairs. As a result, datasets of base-pair and base-stacking interaction were generated, as summarized in **Tables S23** and **S24**.

To evaluate the performance of RNArefine, we used three test datasets designed to assess refinement accuracy under different conditions: (i) benchmark dataset in this study, (ii) Cryo-EM dataset, and (iii) CASP16 dataset.

#### Benchmark dataset in this study

We constructed a benchmark dataset consisting of 63 RNA structures deposited in the Protein Data Bank (PDB) after 2022. These structures are not included in the training datasets of this study. The sequence lengths of the RNA molecules range from 32 to 395 nucleotides.

#### Cryo-EM dataset

For evaluation on Cryo-EM-predicted structures, we used a subset of the EMRNA test dataset. Specifically, among the 71 RNA structures in the original EMRNA main test set, we excluded structures whose RNA sequences were identical to those included in the training data. The resulting Cryo-EM dataset consists of 15 RNA structures. For all targets in this dataset, the initial structures for refinement were obtained from EMRNA models identified directly from Cryo-EM density maps.

#### CASP16 structure dataset

To assess performance under CASP evaluation conditions, we used a subset of monomer targets from CASP16 for which experimentally determined structures are publicly available in the PDB. This dataset consists of 16 RNA monomer targets with the following target IDs: R1203, R1205, R1209, R1211, R1212, R1242, R1251, R1261, R1262, R1263, R1264, R1283, R1285, R1286, R1288, and R1296. This dataset enables direct comparison with CASP16 assessment results and supports CASP-consistent evaluation of refined structures.

### Derivation of Statistical Parameters and Structural Fragments

Using the structures with a resolution of 2.5 Å or better, we calculated statistical parameters for bond lengths, bond angles, torsion angles, and hydrogen bonding. The same dataset was also employed to obtain structure fragments for base-pairs, base-stacking, and single nucleotides.

To generate these fragments, k-means clustering was applied to local nucleotide coordinates. These coordinates were defined using the positions of N9, C4, and C8 atoms for purines and N1, C2, and C6 atoms for pyrimidines, as described by Gendron et al.^28^. For base-pair and base-stacking fragments, the local coordinates of both nucleotides were computed in each of the two local coordinate systems, and the resulting coordinate sets were concatenated. For single nucleotide fragments, clustering was applied to the local coordinate representations of each nucleotide. Based on the structure closest to the center of each cluster, we generated 150, 50, and 150 fragments for base-pairs, base-stacking, and single nucleotide fragments, respectively.

### Prediction of base-pair and base-stacking interactions

To identify base-pair and base-stacking interactions on backbone frames or C4’-traces, neural network models based on geometric attention^39^ were constructed (**Fig. 1B**). In the backbone frame-based prediction, we defined local coordinate frames by constructing two frames using the atom triplets (P, C4′, C3′) and (O5′, C3′, O3′). The final prediction score was obtained by averaging the predictions from these two frames. In contrast, for C4′-trace inputs, where directional information is unavailable, the attention computation relies solely on distance-based geometric features. To further enhance the representation of RNA-specific geometric features, we proposed parameterized geometric normalization into the attention computation.

In this interaction prediction, base-pair interactions were first predicted, and the resulting predictions were then used to predict base-stacking interactions. Base-pair and base-stacking interactions are predicted for all residue pairs (*i, j*) within 20 Å and 15 Å of the C4’ atoms, respectively. In addition to the target residues *i* and *j*, information from neighboring residues in the sequence (*i*±1, *i*±2, *j*±1, *j*±2) is incorporated. For base-stacking prediction, residues base-paired with *i* and *j* and their sequence neighbors are also considered to capture stacking mediated by base-pairing. Furthermore, spatially neighboring residues are added as needed, and the total number of residues considered, including *i* and *j*, is up to 16.

Given an input RNA sequence *X* = (*x*_1_, *x*_2_, *x*_3_, … *x*_*L*_) with a length of *L*, each nucleotide *x*_*k*_ is first mapped to a continuous vector representation *s*_*k*_ ∈ ℝ^*d*^ using a learnable token embedding layer applied to its one-hot representation:

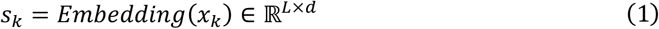

where *s*_*k*_ denotes the token embedding of the *k*-th nucleotide, and *d* is set to 256.

To incorporate sequence-order information relative to the reference residues *i* and *j*, the relative position of each nucleotide *k* is clipped to the range [*-*32, 32] and embedded as vectors 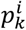 and 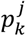 using a relative position embedding layer:

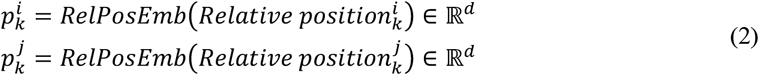

Each of these relative position embeddings is then scaled by a learnable scalar parameter *α* ∈ ℝ, and added to the nucleotide embedding *s*_*k*_ to obtain representations that are aware of the relative positions of residues *i* and *j*, respectively:

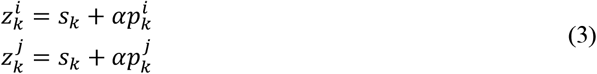

These two representations are concatenated and passed through a linear transformation to produce the position-aware nucleotide representation:

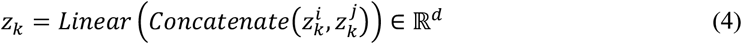

The resulting representation *z*_1_ encodes the relative positional information of residue *k* with respect to residues *i* and *j*. These representations are then further processed through geometric attention layers. Specifically, for each residue *k* the position-aware representation *z*_*k*_ is linearly projected to obtain directional and positional features used to compute the query and key vectors:

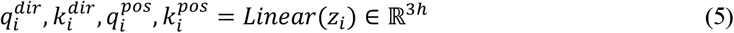

where *h* is the number of attention heads and set to 128. To incorporate structure information including directions and distances between residues, these vectors are transformed into the global coordinate system using the local frame defined by the rotation matrix *Rot*_*i*_ ∈ *SO*(3) and the translation vector *t*_*i*_ ∈ ℝ^3^ associated with each nucleotide *i*. The directional features are first rotated into the global coordinate system using the local rotation matrices:

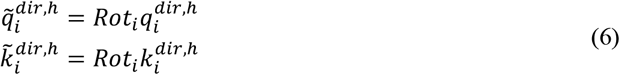

The dot product between these transformed vectors defines the rotation-based term:

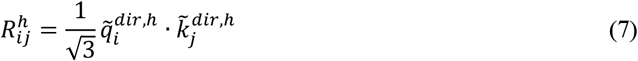

In parallel, the positional features are transformed by both rotation and translation to reflect their absolute spatial positions:

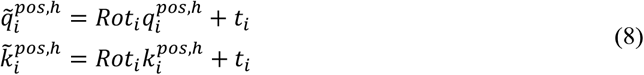

The Euclidean distance between these transformed positions gives the distance-based term:

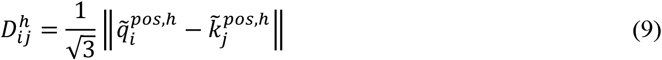

In contrast with the original geometric attention mechanism, to introduce learnable centering and scaling that define head-specific optimal distances and rotations, the distance *D* and direction *R* terms were standardized prior to the computation of attention as follows:

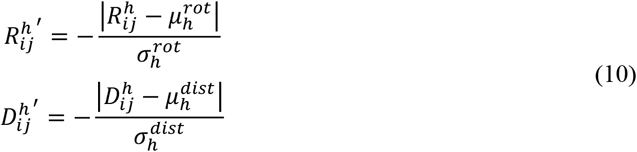

where 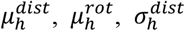 and 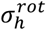 are learnable parameters. In backbone frame-based base-pair prediction, the attention between residues is computed as a weighted combination of the direction and distance terms:

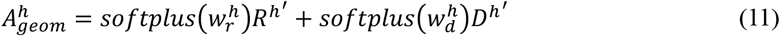

where *w*_*r*_ and *w*_*d*_ are head-specific scalar parameters that control the relative weighting of the rotation and distance terms, respectively. In contrast, in C4′-trace-based prediction, the attention is computed solely from the distance term.

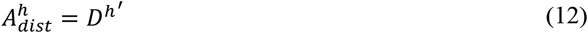

For base-stacking prediction, predicted base-pair information is incorporated into the attention mechanism by projecting the binary base-pair map into a head-specific embedding space using a learnable embedding layer, yielding 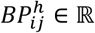, and a base-pair term is added in the calculation of attention as follows:

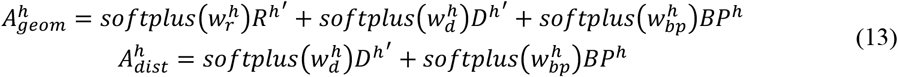

where *w*_*bp*_ is a head-specific scalar parameter for the base-pair term. Using the (*i, j*)-th element of the resulting matrix, denoted as 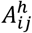, the aggregated representation is obtained by taking the weighted sum of the value vectors 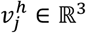:

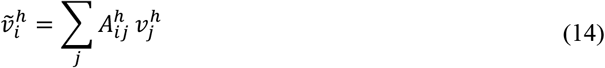

where 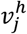 is defined in the global coordinate system by rotating 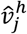, which is a linear projection of the position-aware nucleotide representation, with the rotation matrix 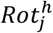:

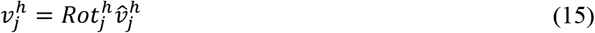

To obtain the weighted-sum vector 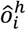 in the local coordinate system, the inverse rotation 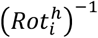 is applied:

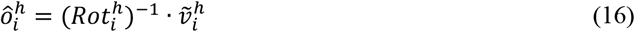

These outputs are concatenated over all heads and then linearly projected:

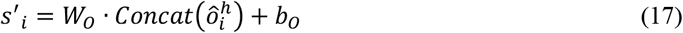

where *W*_*O*_ and *b*_*O*_ are a learnable weight matrix and a learnable bias, respectively. The output *x* is fed to a feed-forward network (FFN) composed of layer normalization, two linear transformations, and a SwiGLU activation as follows.

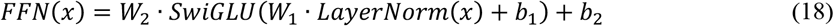

where *W*_1_ and *W*_2_ are learnable weight matrices, and *b*_1_ and *b*_2_ are learnable biases. The hidden dimension after the first linear projection is set to 1024. We stack five blocks, where each block consists of a geometric attention layer and a feed-forward network (FFN). After the final block, the output representations corresponding to the residue *i* and *j* are extracted and concatenated. This vector is then passed through an output layer composed of two linear transformations, where the first linear transformation is followed by layer normalization, GELU activation, and dropout layer with a hidden dimension of 256, and the second linear transformation projects the hidden representation to a scalar score.

The models were trained on the dataset described in **“Dataset construction”**, using 90% of the data for training and 10% for validation. Base-stacking interactions were categorized into two stacking types, adjacent and non-adjacent base-stacking. Separate models were constructed for each nucleotide pair and interaction type (10 base-pair, 16 adjacent base-stacking, and 10 non-adjacent base-stacking). The training was conducted using mini batch learning with a batch size of 64. Binary cross-entropy loss was used for adjacent stacking prediction, while the loss was weighted in base-pair and non-adjacent stacking predictions due to class imbalance:

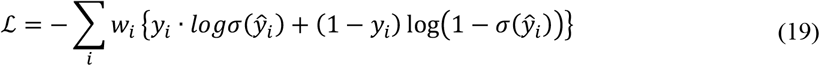

where *y*_*i*_ ∈ {0,1} is the ground truth label indicating whether the residue pair forms a base-pair or stacking interaction (*y*_*i*_=1) or not (*y*_*i*_=0). 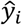 is the predicted logit, *w*_*i*_ is a sample-specific weight, and σ(∗) denotes the sigmoid function. The value of weights *w*_*i*_ is determined by the label *y*_*i*_ as follows:

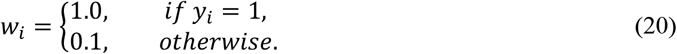

The optimization was performed using the Adam optimizer, with an initial learning rate of 0.00001, and was terminated if the F1-score on the validation data did not improve for 10 consecutive epochs.

For canonical and wobble base pairs, we adopted a postprocessing approach similar to that of Fu et al.^40^, to satisfy the structural constraints of RNA secondary structures. In the postprocessing, base-pair scores below 0.5 were set to zero, then the optimal combination of base-pairs was selected using the SCIP solver so that the total score was maximized under the constraint that each nucleotide forms a base-pair with at most one partner. In contrast to Fu et al., we did not introduce a constraint to exclude sharp loops, since such loops are observed in actual 3D RNA structures.

### Full-atom structure generation

Full-atom structures were generated by superimposing base-pair and single-nucleotide fragments in loop regions onto the backbone structure based on the predicted base-pair information. For nucleotide pairs predicted as Watson–Crick or wobble pairs, we superimposed the corresponding base-pair fragments. In all other cases, single-nucleotide fragments in loop regions were used. The fragment superposition was performed using the Kabsch algorithm^41^, and among the 150 fragments generated via clustering in **“Derivation of Statistical Parameters and Structural Fragments”**, the fragment with the lowest root-mean-square deviation (RMSD) was selected.

#### Structure refinement

To refine structures, two types of refinement methods were applied: MC-based refinement and L-BFGS-based refinement. These refinements were performed based on the predicted base-pair and base-stacking information, and in both refinements all atomic coordinates were adjusted during the process. In these procedures, all parameters required for the energy calculations and structural movements were generated from the high-resolution structures (see **“Derivation of Statistical Parameters and Structural Fragments”**).

#### Monte Carlo simulation-based structure refinement

In the MC-based refinement, Metropolis criterion was applied at a temperature of 3.0 to decide whether each movement was accepted or rejected based on the energy difference before and after the movement. The refinement was performed for up to 50,000 MC steps or until 2 hours of computation time had elapsed.

### Force fields

The energy consists of 10 terms:

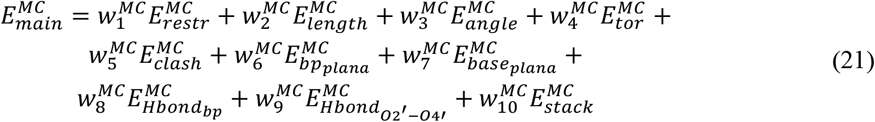

where *w*_*i*_ are weights assigned to each energy term. The weights were empirically determined to balance the contributions of individual energy terms. The values of 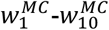 were set to 3.0, 60.0, 1.0, 1.0, 120, 40.0, 35.0, 60.0, 10.0, and 30.0, respectively.

The distance restraint energy term, 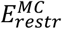 was used to maintain the interatomic distance between two atoms close to their reference structure. It is defined as:

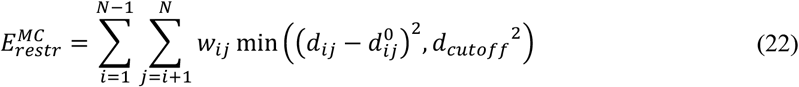

where *N* is the total number of residues in the structure, and *d*_*ij*_ and 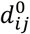 denote the distances between backbone atoms (P and C4’) of residues *i* and *j* in the refining and initial structures, respectively. The cutoff distance *d*_*cutoff*_ was set to 20 Å, limiting the maximum penalty assigned to large deviations. The weighting factor *w*_*ij*_ was introduced to reflect the flexibility of different structural regions, with a weight of 0.1 for restraints involving external loops and 1.0 for all other regions.

The bond length and inner angle energy terms 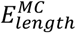 and 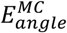 enforce each bond length and angles to remain close to their statistical mean values, respectively. For example, the bond length energy for bond *i* is calculated as:

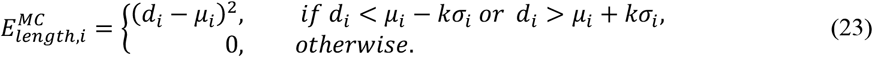

where *d*_*i*_ is the bond length in the refining structure, and *μ*_*i*_ and σ_*i*_ are its mean and standard deviation. *k* is a scaling factor that defines the width of the allowed range, and it was set to 3.5. The total bond length energy was obtained by summing over all bond lengths in the structure. The inner angle restraint energy was calculated in the same manner.

The torsion angle energy term, 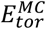 was defined using a potential discretized into 360 bins spanning the full dihedral range (0–360°). For each bin, the statistical frequency was converted into an energy value 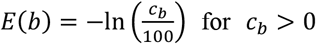, and bins with zero counts were set to *E*(*b*) = 0. The torsion angles in the refining structure were then mapped to their corresponding bin, and the total energy was obtained by summing the energy values assigned to these bins.

To prevent steric overlaps between atoms, a steric clash energy term, 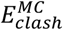, was introduced. If the distance between two atoms, *r*_*ij*_, is closer than the sum of their van der Waals radii, *vdW*(*i*) + *vdW*(*j*), a penalty according to the overlap distance is applied as follows:

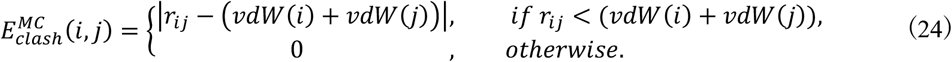

The total clash energy was obtained by summing these penalties.

In canonical and wobble base pairs, the two bases lie nearly on the same plane. To ensure coplanarity of base-pairs, we incorporated the base-pair plane energy term, 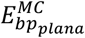, which was originally proposed in QRNAS^22^. In the energy calculation, a best-fit plane is determined so that the total squared perpendicular distance of all atoms from this plane is minimized, and the total squared distance of all base atoms from the plane defines 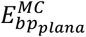. To suppress the influence of false positives and emphasize the potential for high-confidence base-pairs, the base-pair plane energy term for each base-pair was weighted by the predicted base-pair probability. Given the predicted probability *x* ∈ [0,1] for a base-pair, the weight was computed as:

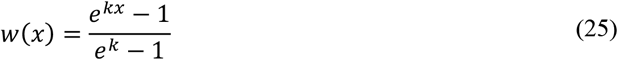

where the scaling parameter was fixed at *k* = 2.0. The weighted plane energies were then summed over all base-pairs. Similar to 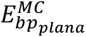, to preserve the planarity of each nucleotide base, a base plane energy term 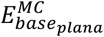 was introduced. Instead of using torsion angles, 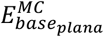 was calculated in the same manner as 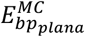, by minimizing the total squared perpendicular distances of atoms in each base from its best-fit plane. To refine the geometry of hydrogen bonding, two hydrogen-bond energy terms, 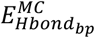 and 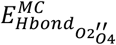 were introduced. These terms regulate the donor–acceptor and hydrogen–acceptor distances, as well as the donor– hydrogen–acceptor angle

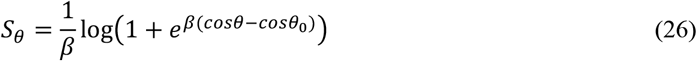

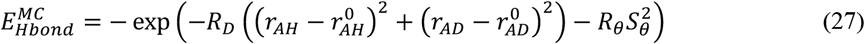

where *r*_*AH*_ and *r*_*AD*_ denote the current acceptor–hydrogen and acceptor–donor distances, and 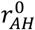 and 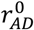 are their statistical values. The angular deviation term *S*_*θ*_ is computed using a smooth one-sided softplus function, where *θ* is the donor–hydrogen–acceptor angle value and *θ*_0_ is its statistical value. The constants *R*_*D*_, *R*_*θ*_, and *β* control stiffness and smoothness of the potential and were set to 2.0, 2.0, and 25, respectively. The 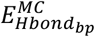 is for hydrogen bonds between base-pairs, including both canonical and non-canonical pairs, while 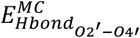 is for hydrogen bonds formed between the O2′ (donor) and O4′ (acceptor) atoms. For canonical base-pairs, hydrogen bonds were identified between donor–acceptor pairs on the Watson–Crick edges. For non-canonical base-pairs and O2′–O4′ pairs, donor–acceptor interactions were considered when their distances were within *μ*_*AD*_ + 4.0σ_*AD*_, where *μ*_*AD*_ and σ_*AD*_ denote the mean and standard deviation of the donor–acceptor distance, respectively. The 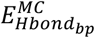 was weighted by the predicted base-pair probability using the same function defined in Eq. 25. These energy terms are knowledge-based potentials inspired by statistically observed geometrical preferences of stable hydrogen bonds, enabling smoother and more reliable geometric refinement.

To form base-stacking interactions, a stacking energy term 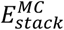 was introduced. This term is used to refine the local stacking geometries. For each base-stacking interaction in the refining structure, stacking fragments were superimposed using the Kabsch algorithm, and the fragment with the smallest RMSD was selected. The stacking energy was then calculated as the sum of deviations between the interatomic distances across the stacked nucleotides in the refining and fragment structure, as follows:

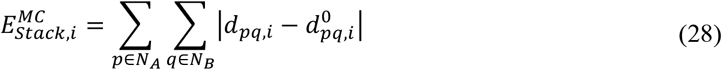

where *d*_*pq*_ and 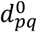 denote the distances between atoms *p* and *q* in the *i*-th stacking interaction of the refining structure and the corresponding fragment, respectively. *N*_*A*_ and *N*_*B*_ represent the numbers of atoms in the stacked nucleotides *A* and *B*. The stacking energy term for each base-stacking was weighted by the predicted stacking probability using the same weighting function defined in Eq. 23.

### Structure movement

To efficiently explore conformational space, the refining structure is represented in two coordinate systems, a length–angle system and a Cartesian coordinate system, and five types of movements are applied. The first three movements modify the length–angle system, in which geometrical parameters including bond lengths, bond angles, and torsion angles are updated, and the Cartesian coordinates of the new structure are subsequently reconstructed from these parameters. In contrast, remaining two movements directly change the atomic positions in Cartesian coordinate system, after which the geometrical parameters are recalculated based on the updated coordinates.

The first move is a bond-length adjustment, which changes the bond-length parameter in two ways. In the first way, the bond length *l* is slightly shifted toward its mean value *μ*, and the step size is adaptively scaled by the deviation from the mean as follows:

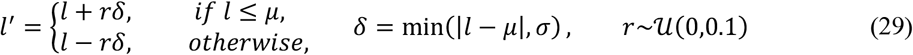

where σ is the standard deviation estimated from the empirical bond-length distribution, and *r* is a random value uniformly drawn between 0 and 0.1. In the second way, the bond length is updated to a random value uniformly sampled from the allowed range [*μ* − 3.5σ, *μ* + 3.5σ].

The second movement is the angle adjustment, which is performed in the same manner as the length movement, except that the updated parameter is the bond angle *θ* instead of the bond length *l*.

The third movement updates torsion angles *ϕ* by sampling from empirical distributions. For each torsion angle type, a histogram is precomputed from experimentally determined RNA structures, covering the range of 0° to 360° with 360 bins of 1° span. In this move, one bin is selected according to its frequency, and then a new torsion value *ϕ*^′^ is uniformly sampled from the range covered by the selected bin.

The fourth movement is a local movement inspired by the LMProt algorithm^42^, which was originally developed for protein structure refinement and was adapted here for RNA. In this movement, a short segment of 2–7 consecutive residues is randomly selected, and the atomic coordinates within the segment are slightly perturbed. The atomic positions are then locally adjusted to satisfy bond-length and bond-angle constraints, ensuring geometrical consistency within the segment.

The last movement is a fragment-based local optimization in Cartesian space. In this movement, a short segment of 2–7 residues is randomly selected, and the coordinates of all residues in the selected fragment and their base-paired residues are refined by an L-BFGS-based optimization process in **“L-BFGS-based structure refinement”** with different energy weighting (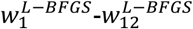 were set to 3.0, 400, 120, 1.0, 1.0, 5.0, 7.5, 17.5, 3.0, 25.0, 7.5, and 0.5 in Equation 30), which is iteratively applied 200 times. Every 25 iterations, for each canonical and wobble base-pair associated with the segment, each base-pair fragment is superimposed onto the corresponding atoms in the refining structure using the Kabsch algorithm, and the base region of the fragment with the lowest RMSD is used to replace the refining base atoms. The optimization was stopped when the total energy, monitored at these 25-step intervals, failed to decrease over three consecutive intervals.

### L-BFGS-based structure refinement

To further refine the structures obtained from the MC-based refinement, we apply a limited-memory BFGS (L-BFGS) optimization with line search. In this refinement, the atomic coordinates are iteratively updated to minimize an energy function composed of twelve differentiable terms. The optimization was terminated when the total energy failed to decrease over 30 steps, or when the maximum runtime of one hour was reached.

The energy consists of 12 terms:

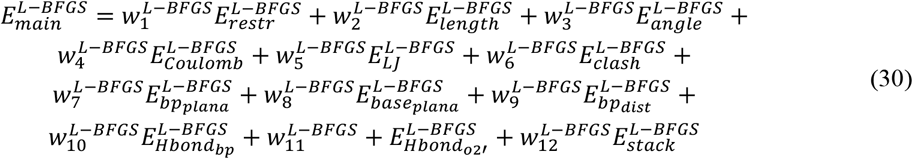

where *w*_*i*_ are empirically determined weights assigned to each term. The values of 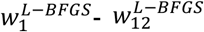 were set to 0.5, 400, 120, 1.0, 1.0, 5.0, 12.5, 30.0, 3.0, 25.0, 7.5, and 0.5, respectively. The 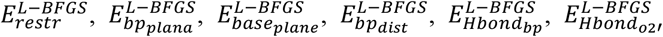 and 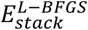 terms are identical to the corresponding MC-based refinement terms 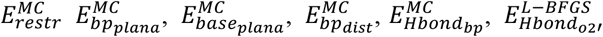, and 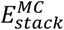 defined in Eq. (21), respectively. The bond-length and bond-angle terms 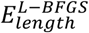 and 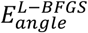 correspond to 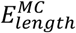 and 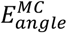 in the MC-based refinement. They are defined as continuous quadratic functions, unlike the conditional form used in MC-based refinement (Eq. 23), enabling smoother gradients for L-BFGS optimization. A base-pair distance restraint energy, denoted as 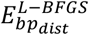, was introduced to stabilize the relative geometry of paired bases. For each base-pair, an optimal base-pair fragment was first selected by superimposing and choosing the one with the minimum RMSD. Then, inter-atomic distances between representative base atoms (N9, C4, C8 for purines; N1, C2, C6 for pyrimidines) were compared between the fragment and the refining structure by penalizing the squared differences of their distances. To capture the fundamental attractive and repulsive forces between atoms, Coulomb and Lennard–Jones potentials, 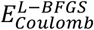 and 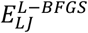, were introduced^43, 44^. The 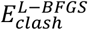 applies a quadratic penalty when the interatomic distance is shorter than the sum of the van der Waals radii.

### Evaluation

#### Evaluation for base-pair and base-stacking prediction

For the evaluation of base-pair and base-stacking prediction, we compared the predicted interactions with ground-truth interactions obtained by applying ClaRNA to native structures. The performance was evaluated by precision, recall, F1-score, and Matthews correlation coefficient (MCC) as follows:

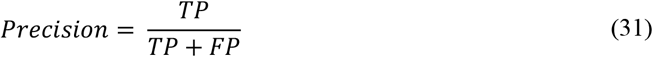

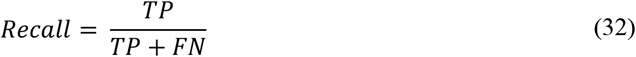

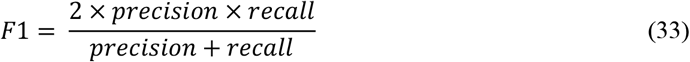

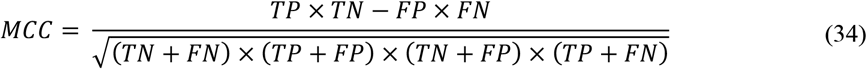

where TP, FP, TN, and FN correspond to true positive, false positive, true negative, and false negative, respectively. In addition to native structures, interaction identification was also evaluated on predicted structures generated by different structure prediction methods. For this evaluation, we used structures generated by two sequence-based RNA 3D structure prediction methods, DRfold2 and AlphaFold3, as well as one Cryo-EM–based method, EMRNA. The evaluation was performed using DRfold2 and AlphaFold3 models on the benchmark dataset in this study, and using EMRNA models on a Cryo-EM dataset. (see **“Dataset construction”**).

#### Evaluation metrics for structure refinement

To comprehensively evaluate the performance of structure refinement, we assessed models using 16 metrics that capture diverse structural aspects, including global topology, local accuracy, geometric validity, and interaction fidelity. Global structural similarity to experimental structures was evaluated using root-mean-square deviation (RMSD), TM-score^32, 45^, and GDT-TS^34^, all of which were computed using the TM-score package. Geometric quality and stereochemical correctness were assessed using Clashscore, RMSD of bond lengths, RMSD of bond angles, and the MolProbity score, all computed with the MolProbity software^46^. Local structural accuracy was evaluated using lDDT (Local Distance Difference Test)^30^, which measures the consistency of local interatomic distances between predicted and experimental structures, as implemented in the OpenStructure toolkit^47^. In addition to the standard lDDT, we also used lDDT scores penalized for steric clashes and chemical violations following the CASP16 evaluation protocol. Similar to penalized lDDT, we introduced a penalized TM-score to account for both global folding and geometric validity. Specifically, we utilized the OpenStructure toolkit to identify severe stereochemical violations like steric clashes, bad bond lengths, and bad bond angles. The penalized TM-score is mathematically defined as follows:

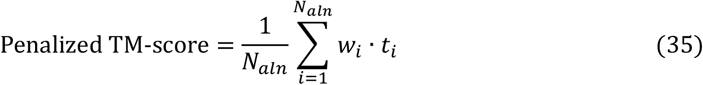

where *N*_*aln*_ is the total number of aligned residues, and *t*_*i*_ is the TM-score of the *i*th residue^32^, *w* = 0 if *i* ∈ *V*, or =1 otherwise. The *V* denotes the set of residues identified as having one or more stereochemical violations. By applying the binary penalty weight *w*_*i*_ to the *i*th residue, this formulation ensures that any residue involved in physically implausible geometries is strictly excluded from contributing to the final global score.

To assess the accuracy of base-pair and base-stacking interactions, interaction network fidelity (INF)^48^ metrics were calculated using rna-tools. We report INF_all, which considers all interactions, as well as interaction-specific metrics including INF_stack (base stacking), INF_wc (Watson–Crick base pairs), and INF_nwc (non–Watson–Crick base pairs). Structural similarity in torsion angle space was evaluated using Mean of Circular Quantities (MCQ)^49^, providing a complementary, sequence-independent measure of conformational similarity. To evaluate hydrogen-bonding accuracy at the atomic level, including both base-pairing and non–base-pairing hydrogen bonds, we employed the HB-score^50^, defined as the ratio of correctly predicted hydrogen bonds to the total number of hydrogen bonds in the experimental structure. Hydrogen bonds were identified using HBPLUS^51^.

In addition to individual metrics, we defined a composite score to measure the structure quality using a single scalar value. For each target, individual evaluation metrics were standardized across different models by converting raw metric values into Z-scores. Prior to standardization, outlier values exceeding ±2 standard deviations from the mean were excluded to reduce the influence of extreme values. For metrics where lower values indicate better structural quality (e.g., RMSD, Clashscore, and geometric deviations), metric values were sign-inverted so that higher Z-scores consistently correspond to better performance. To minimize redundancy among related metrics, the 14 selected metrics were grouped into four categories: global fold similarity (TM-score, GDT-TS, and RMSD), interaction fidelity (INF_stack, INF_wc, and INF_nwc), stereochemical validity (MolProbity score, clashscore, RMSD of bond lengths, RMSD of bond angles, and HB-score), and physically penalized structural quality (penalized TM-score and penalized lDDT). The Z-scores were then averaged within each category, and the resulting category scores were summed across all categories as follows:

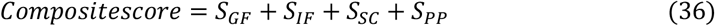

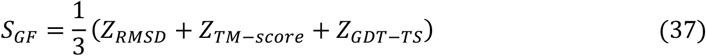

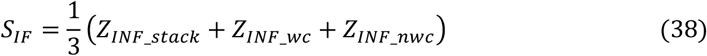

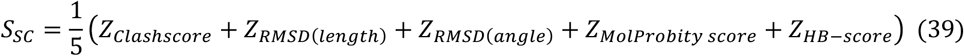

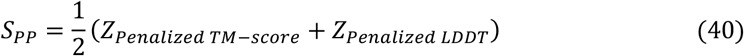

where each *Z*_∗_ denotes the Z-score of the corresponding evaluation metric described above. Notably, to avoid bias from unequal interaction frequencies, we used INF_stack, INF_wc, and INF_nwc separately rather than an aggregated INF_all score.

#### Setting of existing refinement methods

We compared the performance of RNArefine with that of PDBfixer, RNAfitme, Arena, QRNAS, and Molecular dynamics (MD) simulation. PDBfixer was executed using its default parameters with the publicly available source code. RNAfitme was run under standard settings through its public web server. Arena offers two operational modes. One mode preserves the input atomic coordinates, while the other updates them during refinement. To enable full structural optimization rather than restricting adjustments to added atoms only, we executed the latter mode. QRNAS requires substantial computational time when executed with its default settings. To ensure a fair comparison under same computational conditions, we limited its runtime to three hours, which matches the maximum simulation time used in our refinement. Molecular dynamics simulations were conducted using the same protocol and parameter settings as those described in the original RhoFold study. Since PDBfixer and Arena are capable of generating full-atom structures from coarse-grained models, missing atoms were complemented according to each method’s reconstruction procedure. For the other refinement methods (RNAfitme, QRNAS, and MD simulation), the same initial full-atom structures used for RNArefine were refined.

### CASP score and ranking estimation of refined structure

To assess the refinement capability of RNArefine on diverse initial structures, we applied it to models submitted to CASP16. For evaluation purposes, we focused on 16 targets whose experimentally determined structures are available in the PDB. For each of these targets, refinement was applied to the first models submitted by the top 30 groups based on their performance on these 16 targets. Prior to refinement, base-pair and base-stacking interactions were predicted from the backbone frames of these models and subsequently used as structural restraints during refinement.

CASP is conducted as a strictly blind experiment, and while predicted models are released after the competition, the native structures used for official evaluation are not publicly available. For some targets, experimentally determined structures are later deposited in the Protein Data Bank (PDB); however, these structures are not guaranteed to be identical to the CASP native structures. Therefore, evaluation scores computed directly against PDB references should not be compared with CASP scores. In this study, to enable comparison of structural quality before and after refinement under the CASP evaluation scheme, CASP scores were estimated from the PDB-based performance.

To estimate CASP-equivalent performance for refined models, we focused on TM-score, GDT-TS, and penalized lDDT, which are monomer structural similarity metrics used in CASP assessments. For each target and method, both the original submitted model (decoy) and the refined model were evaluated against the corresponding PDB reference structure. The CASP-equivalent score of the refined model 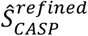 was estimated as:

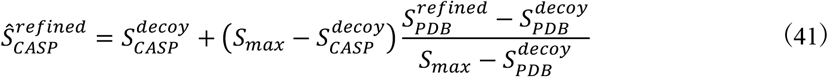

where 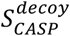 is the official CASP score of the originally submitted model, 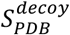 and 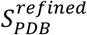 are the scores of the decoy and refined models computed against the PDB reference structure, respectively, and *S*_*max*_ = 1.0 denotes the theoretical maximum of the metric (TM-score, GDT-TS, and penalized lDDT). The values of metrics in the original submitted model in CASP16 were obtained from the study reporting the RNA structure prediction assessment in CASP16^31^. Using the estimated CASP-equivalent scores, refined models were ranked using the same procedure as in the CASP16 assessment. The final evaluation metric, the Z-score, was computed following the CASP16 ranking methodology^31^. The Z-scores for each evaluation metric were then weighted across metrics and summed over all targets following the CASP16 protocol. Because R1261 and R1262, as well as R1263 and R1264, share highly similar sequences, we followed the CASP16 procedure by grouping them together and considering only the higher score between the two. This procedure ensures a fair and CASP-consistent comparison of refined models without requiring access to the original CASP native structures.

## ACKNOWLEDGEMENTS

This work was supported in part by JSPS KAKENHI (25K24407 and 26K21372); the World Research Hub (WRH) Program of Institute of Science Tokyo; the Ministry of Education, Singapore (MOE-T1251RES2309, MOE-T2EP20125-0014, and MOE-T2EP20225-0028); the Agency for Science, Technology and Research (A*STAR), Singapore (IAF-PP H25J6a0034); and the National Research Foundation, Singapore (NRF-CRP33-2025-0003 and AI4SCT_2025-0014).

## AUTHOR CONTRIBUTIONS

Y.Z. conceived and designed the study framework. S.T., Y.L., K.S., and Y.Z. developed the refinement strategy and designed the statistical potentials and interaction-guided constraints. S.T. implemented the RNArefine pipeline, developed the web server, and carried out benchmarking and validation analyses. Y.L., H.K., and Y.Z. supervised the project and provided scientific guidance. K.S., H.K. and Y.Z. acquired funding for the project. S.T. and Y.Z. wrote the manuscript. All authors reviewed and approved the manuscript.

## DATA AVAILABILITY

To facilitate reproducibility and further methodological development in RNA structure prediction and refinement, we provide the fragment structures, the statistical parameters used in the simulations, and the benchmark dataset constructed in this study on the RNArefine web server (https://zhanggroup.org/RNArefine/). The training data for the base-pair and base-stacking prediction models were obtained from the DRfold^11^ study and PDB^38^. The PDB reference structures used for CASP16 evaluation are publicly available, and the official CASP16 assessment results can be obtained from the CASP16 assessment publication^31^.

## CODE AVAILABILITY

The online server and standalone package of RNArefine are freely available at (https://zhanggroup.org/RNArefine/), where the penalized TM-score package can also be downloaded. Structures were visualized by Pymol v.3.1.0 and ChimeraX v.1.11.1.

## COMPUTATIONAL RESOURCES

Computations were performed in part on the TSUBAME 4.0 supercomputer at Tokyo Institute of Technology, consisting of HPE Cray XD665 compute nodes with dual AMD EPYC 9654 CPUs (192 cores, 768 GiB RAM) and 4 NVIDIA H100 GPUs per node as the base configuration. For base-pair and base-stacking predictions, we used TSUBAME’s node_f resource type (192 CPU cores, 768 GiB RAM, 4 GPUs). For the refinement stage, we used the cpu_16 resource type (16 CPU cores, 36.8 GiB RAM, no GPUs).

