## Supplementary Information for "RNArefine: AI-guided Atomic-Level Refinement of RNA Structures"

Table S15 Comparison of refinement performance among the DRfold2 coarse-grained structure, the initial full-atom structure, refinement using Monte Carlo (MC) simulation only,

### Supplementary Tables

**Table S1. Clashscore, chemical violations, and MolProbity scores for native and AlphaFold-predicted structures in the benchmark dataset used in this study.**

| | Clashscore | RMSD (bond length) | RMSD (bond angle) | MolProbity score |
|---|---|---|---|---|
| **Native** | 5.40 | 0.00514 | 0.903 | 2.27 |
| **AlphaFold3** | 16.2 | 0.0172 | 1.18 | 2.78 |

**Table S2. Clashscore, chemical violations, and MolProbity scores for native and EMRNA-predicted structures in the Cryo-EM dataset.**

| | Clashscore | RMSD (bond length) | RMSD (bond angle) | MolProbity score |
|---|---|---|---|---|
| **Native** | 4.59 | 0.00477 | 0.762 | 2.28 |
| **EMRNA** | 176 | 0.0969 | 10.0 | 3.84 |

**Table S3. Performance comparison of base-pair prediction for native structures in our benchmark dataset.** Sensitivity (SN), Precision (PR), F1-score (F1), and Matthews correlation coefficient (MCC) are reported for all base-pairs, as well as for canonical (Ca) and non-canonical (Nc) base pairs. Evaluations were conducted on the benchmark dataset in this study, which consists of 63 RNA structures.

| | SN | PR | F1 | MCC | SN-Ca | PR-Ca | F1-Ca | MCC-Ca | SN-Nc | PR-Nc | F1-Nc | MCC-Nc |
|---|---|---|---|---|---|---|---|---|---|---|---|---|
| **RNArefine (C4'-trace)** | 0.914 (0.0897) | 0.784 (0.155) | 0.834 (0.124) | 0.808 (0.182) | 0.950 (0.140) | 0.889 (0.141) | 0.916 (0.133) | 0.884 (0.207) | 0.660 (0.303) | 0.465 (0.266) | 0.525 (0.263) | 0.510 (0.264) |
| **RNArefine (Backbone)** | 0.939 (0.100) | 0.856 (0.114) | 0.890 (0.0933) | 0.861 (0.179) | **0.966** (0.0812) | 0.934 (0.0668) | 0.947 (0.0613) | 0.915 (0.176) | **0.817** (0.252) | 0.645 (0.250) | 0.697 (0.227) | 0.681 (0.248) |
| **CSSR** | **0.946** (0.152) | 0.701 (0.194) | 0.790 (0.168) | 0.772 (0.207) | 0.950 (0.150) | 0.864 (0.147) | 0.900 (0.138) | 0.870 (0.208) | 0.724 (0.411) | 0.287 (0.243) | 0.380 (0.271) | 0.401 (0.266) |
| **RNAView** | 0.810 (0.104) | 0.960 (0.0618) | 0.876 (0.0819) | 0.847 (0.171) | 0.923 (0.0871) | 0.976 (0.0541) | 0.948 (0.0682) | 0.916 (0.179) | 0.588 (0.212) | 0.897 (0.178) | 0.698 (0.197) | 0.687 (0.219) |
| **DSSR** | 0.888 (0.100) | **0.982** (0.0319) | **0.930** (0.0683) | **0.900** (0.175) | 0.949 (0.0931) | **0.996** (0.0134) | **0.969** (0.0584) | **0.938** (0.178) | 0.734 (0.187) | **0.947** (0.0961) | **0.816** (0.144) | **0.795** (0.194) |
| **MC-Annotate** | 0.790 (0.139) | 0.920 (0.141) | 0.847 (0.133) | 0.834 (0.168) | 0.932 (0.0742) | 0.969 (0.0617) | 0.948 (0.0574) | 0.916 (0.175) | 0.541 (0.207) | 0.801 (0.226) | 0.632 (0.206) | 0.633 (0.212) |

**Table S4. Performance comparison of base-stacking prediction for native structures in our benchmark dataset.** Sensitivity (SN), Precision (PR), F1-score (F1), and Matthews correlation coefficient (MCC) are reported for all stacking-pairs, as well as for adjacent (Adj) and non-adjacent (NAdj) stacking pairs. Evaluations were conducted on the benchmark dataset in this study, which consists of 63 RNA structures.

| | SN | PR | F1 | MCC | SN-Adj | PR-Adj | F1-Adj | MCC-Adj | SN-NAdj | PR-NAdj | F1-NAdj | MCC-NAdj |
|---|---|---|---|---|---|---|---|---|---|---|---|---|
| **RNArefine (C4'-trace)** | 0.802 (0.0649) | 0.816 (0.0841) | 0.806 (0.0600) | 0.802 (0.0605) | 0.863 (0.0642) | 0.873 (0.0553) | 0.867 (0.0481) | 0.533 (0.171) | 0.596 (0.166) | 0.641 (0.213) | 0.598 (0.157) | 0.590 (0.165) |
| **RNArefine (Backbone)** | 0.899 (0.0450) | **0.894** (0.0552) | **0.895** (0.0388) | 0.893 (0.0394) | 0.937 (0.0440) | **0.949** (0.0278) | **0.943** (0.0281) | 0.792 (0.142) | 0.735 (0.172) | **0.723** (0.195) | 0.716 (0.162) | 0.705 (0.180) |
| **DSSR** | 0.951 (0.0536) | 0.401 (0.159) | 0.549 (0.145) | 0.603 (0.122) | 0.964 (0.0432) | 0.399 (0.166) | 0.547 (0.152) | 0.372 (0.159) | 0.838 (0.278) | 0.368 (0.209) | 0.494 (0.227) | 0.527 (0.224) |
| **MC-Annotate** | **0.993** (0.0129) | 0.812 (0.0611) | 0.892 (0.0372) | **0.895** (0.0349) | **0.992** (0.0166) | 0.872 (0.0563) | 0.927 (0.0338) | **0.801** (0.0825) | **0.996** (0.0160) | 0.611 (0.161) | **0.745** (0.126) | **0.756** (0.139) |

**Table S5. Performance comparison of base-pair prediction for native structures in the Cryo-EM dataset.** Sensitivity (SN), Precision (PR), F1-score (F1), and Matthews correlation coefficient (MCC) are reported for all base-pairs, as well as for canonical (Ca) and non-canonical (Nc) base pairs. Evaluations were conducted on the benchmark dataset in this study, which consists of 15 RNA structures.

| | SN | PR | F1 | MCC | SN-Ca | PR-Ca | F1-Ca | MCC-Ca | SN-Nc | PR-Nc | F1-Nc | MCC-Nc |
|---|---|---|---|---|---|---|---|---|---|---|---|---|
| **RNArefine (C4'-trace)** | 0.914 (0.0617) | 0.868 (0.0977) | 0.887 (0.0635) | 0.888 (0.0624) | 0.948 (0.0513) | 0.953 (0.0482) | 0.949 (0.0341) | 0.948 (0.0341) | 0.636 (0.324) | 0.586 (0.350) | 0.562 (0.296) | 0.584 (0.292) |
| **RNArefine (Backbone)** | 0.922 (0.0545) | 0.914 (0.0646) | 0.917 (0.0532) | 0.917 (0.0536) | 0.954 (0.0567) | 0.976 (0.0308) | **0.964** (0.0351) | **0.963** (0.0352) | **0.766** (0.252) | 0.656 (0.302) | 0.688 (0.262) | 0.699 (0.259) |
| **CSSR** | **0.970** (0.0429) | 0.754 (0.0841) | 0.846 (0.0558) | 0.853 (0.0516) | **0.972** (0.0374) | 0.896 (0.0700) | 0.931 (0.0459) | 0.931 (0.0462) | 0.733 (0.442) | 0.175 (0.147) | 0.273 (0.208) | 0.347 (0.233) |
| **RNAView** | 0.737 (0.0662) | 0.967 (0.0529) | 0.834 (0.0407) | 0.842 (0.0370) | 0.895 (0.0632) | 0.974 (0.0545) | 0.931 (0.0467) | 0.931 (0.0471) | 0.433 (0.185) | 0.929 (0.140) | 0.564 (0.174) | 0.615 (0.147) |
| **DSSR** | 0.865 (0.0962) | **0.989** (0.0240) | **0.921** (0.0635) | **0.923** (0.0607) | 0.934 (0.0641) | **0.994** (0.0148) | 0.963 (0.0399) | 0.962 (0.0399) | 0.669 (0.231) | **0.967** (0.0850) | **0.768** (0.194) | **0.790** (0.168) |
| **MC-Annotate** | 0.737 (0.104) | 0.947 (0.0451) | 0.824 (0.0591) | 0.832 (0.0544) | 0.902 (0.0610) | 0.969 (0.0459) | 0.932 (0.0318) | 0.932 (0.0318) | 0.434 (0.213) | 0.881 (0.163) | 0.549 (0.196) | 0.595 (0.169) |

**Table S6. Performance comparison of base-stacking prediction for native structures in the Cryo-EM dataset.** Sensitivity (SN), Precision (PR), F1-score (F1), and Matthews correlation coefficient (MCC) are reported for all stacking-pairs, as well as for adjacent (Adj) and non-adjacent (NAdj) stacking pairs. Evaluations were conducted on the Cryo-EM dataset in this study, which consists of 15 RNA structures.

| | SN | PR | F1 | MCC | SN-Adj | PR-Adj | F1-Adj | MCC-Adj | SN-NAdj | PR-NAdj | F1-Adj | MCC-NAdj |
|---|---|---|---|---|---|---|---|---|---|---|---|---|
| **RNArefine (C4'-trace)** | 0.836<br>(0.0697) | 0.865<br>(0.0618) | 0.849<br>(0.0607) | 0.846<br>(0.0617) | 0.874<br>(0.0777) | 0.882<br>(0.0475) | 0.877<br>(0.0574) | 0.629<br>(0.170) | 0.710<br>(0.165) | 0.815<br>(0.154) | 0.735<br>(0.127) | 0.747<br>(0.112) |
| **RNArefine (Backbone)** | 0.893<br>(0.0496) | **0.926**<br>(0.0536) | **0.909**<br>(0.0475) | **0.907**<br>(0.0482) | 0.928<br>(0.0550) | **0.952**<br>(0.0301) | **0.940**<br>(0.0405) | **0.828**<br>(0.0989) | 0.758<br>(0.169) | **0.849**<br>(0.160) | **0.783**<br>(0.157) | **0.791**<br>(0.141) |
| **DSSR** | 0.933<br>(0.0603) | 0.500<br>(0.120) | 0.641<br>(0.0914) | 0.673<br>(0.0761) | 0.938<br>(0.0579) | 0.489<br>(0.157) | 0.626<br>(0.125) | 0.426<br>(0.122) | 0.875<br>(0.265) | 0.446<br>(0.180) | 0.580<br>(0.210) | 0.617<br>(0.208) |
| **MC-Annotate** | **0.994**<br>(0.0139) | 0.812<br>(0.0548) | 0.893<br>(0.0369) | 0.896<br>(0.0349) | **0.993**<br>(0.0156) | 0.853<br>(0.0635) | 0.917<br>(0.0418) | 0.810<br>(0.0571) | **0.933**<br>(0.249) | 0.626<br>(0.202) | 0.744<br>(0.216) | 0.761<br>(0.215) |

**Table S7. Performance comparison of base-pair prediction for structures predicted by DRfold2 in our benchmark dataset.** Sensitivity (SN), Precision (PR), F1-score (F1), and Matthews correlation coefficient (MCC) are reported for all base-pairs, as well as for canonical (Ca) and non-canonical (Nc) base pairs. Evaluations were conducted on the benchmark dataset in this study, which consists of 63 RNA structures.

| | SN | PR | F1 | MCC | SN-Ca | PR-Ca | F1-Ca | MCC-Ca | SN-Nc | PR-Nc | F1-Nc | MCC-Nc |
|---|---|---|---|---|---|---|---|---|---|---|---|---|
| **RNArefine (C4'-trace)** | 0.766<br>(0.217) | 0.678<br>(0.236) | 0.709<br>(0.220) | 0.697<br>(0.232) | 0.834<br>(0.226) | 0.785<br>(0.229) | 0.800<br>(0.221) | 0.784<br>(0.241) | 0.517<br>(0.321) | 0.403<br>(0.282) | 0.423<br>(0.265) | 0.422<br>(0.262) |
| **RNArefine (Backbone)** | 0.776<br>(0.222) | **0.700**<br>(0.235) | **0.725**<br>(0.221) | **0.713**<br>(0.233) | 0.833<br>(0.225) | **0.791**<br>(0.228) | **0.803**<br>(0.221) | **0.787**<br>(0.241) | 0.592<br>(0.311) | **0.491**<br>(0.287) | **0.499**<br>(0.259) | **0.500**<br>(0.258) |
| **CSSR** | **0.835**<br>(0.228) | 0.618<br>(0.225) | 0.694<br>(0.217) | 0.691<br>(0.225) | **0.843**<br>(0.224) | 0.768<br>(0.222) | 0.794<br>(0.215) | 0.779<br>(0.235) | **0.607**<br>(0.424) | 0.243<br>(0.225) | 0.322<br>(0.259) | 0.350<br>(0.263) |

**Table S8. Performance comparison of base-stacking prediction for structures predicted by DRfold2 in our benchmark dataset.** Sensitivity (SN), Precision (PR), F1-score (F1), and Matthews correlation coefficient (MCC) are reported for all stacking-pairs, as well as for adjacent (Adj) and non-adjacent (NAdj) stacking pairs. Evaluations were conducted on the benchmark dataset in this study, which consists of 63 RNA structures.

| | SN | PR | F1 | MCC | SN-Adj | PR-Adj | F1-Adj | MCC-Adj | SN-NAdj | PR-NAdj | F1-NAdj | MCC-NAdj |
|---|---|---|---|---|---|---|---|---|---|---|---|---|
| **RNArefine (C4'-trace)** | 0.743<br>(0.0915) | 0.763<br>(0.118) | 0.748<br>(0.0896) | 0.744<br>(0.0901) | **0.814**<br>(0.0878) | 0.839<br>(0.0874) | 0.823<br>(0.0709) | 0.388<br>(0.193) | 0.486<br>(0.197) | 0.527<br>(0.258) | 0.479<br>(0.195) | 0.489<br>(0.195) |
| **RNArefine (Backbone)** | **0.755**<br>(0.0955) | **0.824**<br>(0.111) | **0.784**<br>(0.0912) | **0.781**<br>(0.0925) | 0.799<br>(0.0930) | **0.914**<br>(0.0738) | **0.849**<br>(0.0718) | **0.416**<br>(0.231) | **0.544**<br>(0.223) | **0.546**<br>(0.261) | **0.521**<br>(0.220) | **0.529**<br>(0.216) |

**Table S9. Performance comparison of base-pair prediction for structures predicted by AlphaFold3 in our benchmark dataset.** Sensitivity (SN), Precision (PR), F1-score (F1), and Matthews correlation coefficient (MCC) are reported for all base-pairs, as well as for canonical (Ca) and non-canonical (Nc) base pairs. Evaluations were conducted on the benchmark dataset in this study, which consists of 63 RNA structures.

| | SN | PR | F1 | MCC | SN-Ca | PR-Ca | F1-Ca | MCC-Ca | SN-Nc | PR-Nc | F1-Nc | MCC-Nc |
|---|---|---|---|---|---|---|---|---|---|---|---|---|
| **RNArefine (C4'-trace)** | 0.762<br>(0.261) | 0.686<br>(0.265) | 0.710<br>(0.252) | 0.714<br>(0.251) | 0.815<br>(0.270) | 0.808<br>(0.258) | 0.804<br>(0.257) | 0.803<br>(0.258) | 0.522<br>(0.318) | 0.396<br>(0.297) | 0.417<br>(0.266) | 0.434<br>(0.269) |
| **RNArefine (Backbone)** | 0.765<br>(0.263) | 0.717<br>(0.277) | **0.724**<br>(0.258) | **0.729**<br>(0.255) | 0.819<br>(0.275) | 0.822<br>(0.262) | 0.813<br>(0.262) | 0.812<br>(0.264) | 0.558<br>(0.309) | 0.493<br>(0.309) | **0.486**<br>(0.278) | **0.502**<br>(0.276) |
| **CSSR** | **0.815**<br>(0.269) | 0.639<br>(0.249) | 0.700<br>(0.245) | 0.710<br>(0.242) | **0.823**<br>(0.273) | 0.795<br>(0.253) | 0.801<br>(0.255) | 0.800<br>(0.257) | **0.639**<br>(0.407) | 0.280<br>(0.248) | 0.363<br>(0.276) | 0.373<br>(0.271) |
| **ClaRNA** | 0.756<br>(0.259) | 0.721<br>(0.271) | 0.723<br>(0.256) | 0.728<br>(0.253) | 0.822<br>(0.269) | 0.842<br>(0.242) | **0.818**<br>(0.256) | **0.820**<br>(0.251) | 0.522<br>(0.324) | 0.477<br>(0.333) | 0.467<br>(0.306) | 0.479<br>(0.305) |
| **RNAView** | 0.619<br>(0.222) | **0.749**<br>(0.266) | 0.669<br>(0.235) | 0.672<br>(0.235) | 0.776<br>(0.257) | **0.845**<br>(0.242) | 0.797<br>(0.250) | 0.797<br>(0.246) | 0.337<br>(0.199) | **0.574**<br>(0.310) | 0.406<br>(0.219) | 0.426<br>(0.226) |
| **DSSR** | 0.671<br>(0.241) | 0.747<br>(0.268) | 0.698<br>(0.248) | 0.700<br>(0.248) | 0.795<br>(0.263) | 0.844<br>(0.243) | 0.806<br>(0.254) | 0.806<br>(0.250) | 0.418<br>(0.244) | 0.571<br>(0.313) | 0.459<br>(0.255) | 0.473<br>(0.257) |
| **MC-Annotate** | 0.629<br>(0.221) | 0.734<br>(0.269) | 0.667<br>(0.233) | 0.671<br>(0.234) | 0.786<br>(0.259) | 0.842<br>(0.242) | 0.800<br>(0.250) | 0.801<br>(0.246) | 0.329<br>(0.180) | 0.522<br>(0.306) | 0.382<br>(0.210) | 0.400<br>(0.217) |

**Table S10. Performance comparison of base-stacking prediction for structures predicted by AlphaFold3 in our benchmark dataset.** Sensitivity (SN), Precision (PR), F1-score (F1), and Matthews correlation coefficient (MCC) are reported for all stacking-pairs, as well as for adjacent (Adj) and non-adjacent (NAdj) stacking pairs. Evaluations were conducted on the benchmark dataset in this study, which consists of 63 RNA structures.

| | SN | PR | F1 | MCC | SN-Adj | PR-Adj | F1-Adj | MCC-Adj | SN-NAdj | PR-NAdj | F1-Adj | MCC-NAdj |
|---|---|---|---|---|---|---|---|---|---|---|---|---|
| RNArefine (C4'-trace) | 0.724 (0.0930) | 0.794 (0.104) | 0.754 (0.0876) | 0.750 (0.0902) | 0.787 (0.0934) | 0.879 (0.0640) | 0.827 (0.0682) | 0.342 (0.213) | 0.486 (0.206) | 0.537 (0.258) | 0.483 (0.201) | 0.493 (0.200) |
| RNArefine (Backbone) | 0.750 (0.102) | 0.834 (0.104) | 0.786 (0.0942) | 0.783 (0.0967) | 0.796 (0.100) | 0.926 (0.0457) | 0.853 (0.0706) | 0.403 (0.239) | 0.545 (0.229) | **0.544** (0.283) | 0.512 (0.226) | 0.524 (0.223) |
| ClaRNA | 0.767 (0.105) | **0.847** (0.101) | **0.802** (0.0959) | **0.799** (0.0983) | 0.799 (0.102) | **0.944** (0.0467) | **0.863** (0.0737) | **0.442** (0.246) | 0.603 (0.245) | 0.534 (0.263) | **0.542** (0.233) | **0.552** (0.231) |
| DSSR | 0.672 (0.161) | 0.289 (0.121) | 0.394 (0.121) | 0.425 (0.120) | 0.699 (0.156) | 0.325 (0.134) | 0.428 (0.129) | -0.0195 (0.194) | 0.460 (0.357) | 0.145 (0.138) | 0.210 (0.188) | 0.247 (0.204) |
| MC-Annotate | **0.803** (0.111) | 0.779 (0.0862) | 0.788 (0.0887) | 0.784 (0.0919) | **0.812** (0.105) | 0.899 (0.0460) | 0.850 (0.0687) | 0.416 (0.250) | **0.725** (0.281) | 0.396 (0.222) | 0.488 (0.227) | 0.520 (0.225) |

**Table S11. Performance comparison of base-pair prediction for structures predicted by EMRNA in the Cryo-EM dataset.** Sensitivity (SN), Precision (PR), F1-score (F1), and Matthews correlation coefficient (MCC) are reported for all base-pairs, as well as for canonical (Ca) and non-canonical (Nc) base pairs. Evaluations were conducted on the Cryo-EM–determined structure dataset, which consists of 15 RNA structures.

| | SN | PR | F1 | MCC | SN-Ca | PR-Ca | F1-Ca | MCC-Ca | SN-Nc | PR-Nc | F1-Nc | MCC-Nc |
|---|---|---|---|---|---|---|---|---|---|---|---|---|
| RNArefine (C4'-trace) | **0.812** (0.157) | **0.695** (0.186) | **0.745** (0.169) | **0.748** (0.168) | **0.872** (0.126) | **0.801** (0.182) | **0.832** (0.159) | **0.831** (0.159) | 0.399 (0.367) | **0.301** (0.326) | **0.325** (0.325) | **0.336** (0.328) |
| RNArefine (Backbone) | 0.768 (0.168) | 0.576 (0.190) | 0.653 (0.181) | 0.660 (0.178) | 0.786 (0.163) | 0.699 (0.220) | 0.733 (0.192) | 0.732 (0.192) | 0.290 (0.380) | 0.0952 (0.130) | 0.139 (0.184) | 0.162 (0.211) |
| CSSR | 0.800 (0.130) | 0.630 (0.159) | 0.701 (0.146) | 0.706 (0.143) | 0.821 (0.127) | 0.754 (0.176) | 0.782 (0.151) | 0.780 (0.151) | **0.450** (0.449) | 0.135 (0.153) | 0.199 (0.213) | 0.238 (0.242) |
| ClaRNA | 0.704 (0.161) | 0.584 (0.191) | 0.631 (0.171) | 0.635 (0.170) | 0.745 (0.150) | 0.692 (0.221) | 0.711 (0.185) | 0.708 (0.186) | 0.301 (0.370) | 0.167 (0.259) | 0.201 (0.275) | 0.215 (0.283) |
| RNAView | 0.513 (0.161) | 0.636 (0.169) | 0.557 (0.142) | 0.562 (0.143) | 0.662 (0.143) | 0.738 (0.196) | 0.689 (0.152) | 0.688 (0.155) | 0.150 (0.185) | 0.223 (0.280) | 0.165 (0.193) | 0.173 (0.204) |
| DSSR | 0.583 (0.184) | 0.638 (0.186) | 0.606 (0.180) | 0.605 (0.181) | 0.712 (0.163) | 0.733 (0.200) | 0.720 (0.179) | 0.715 (0.182) | 0.212 (0.240) | 0.259 (0.277) | 0.217 (0.229) | 0.225 (0.237) |
| MC-Annotate | 0.271 (0.150) | 0.633 (0.152) | 0.364 (0.154) | 0.397 (0.144) | 0.492 (0.169) | 0.732 (0.190) | 0.576 (0.159) | 0.582 (0.162) | 0.0564 (0.0848) | 0.211 (0.199) | 0.0823 (0.111) | 0.0984 (0.120) |

**Table S12. Performance comparison of base-stacking prediction for structures predicted by EMRNA in the Cryo-EM dataset.** Sensitivity (SN), Precision (PR), F1-score (F1), and Matthews correlation coefficient (MCC) are reported for all stacking-pairs, as well as for adjacent (Adj) and non-adjacent (NAdj) stacking pairs. Evaluations were conducted on the Cryo-EM dataset in this study, which consists of 15 RNA structures.

| | SN | PR | F1 | MCC | SN-Adj | PR-Adj | F1-Adj | MCC-Adj | SN-NAdj | PR-NAdj | F1-Adj | MCC-NAdj |
|---|---|---|---|---|---|---|---|---|---|---|---|---|
| RNArefine (C4'-trace) | 0.796 (0.0771) | **0.731** (0.126) | **0.759** (0.102) | **0.756** (0.101) | 0.849 (0.0742) | **0.781** (0.0876) | **0.811** (0.0730) | **0.486** (0.179) | 0.613 (0.202) | **0.600** (0.274) | **0.578** (0.211) | **0.590** (0.207) |
| RNArefine (Backbone) | 0.814 (0.0889) | 0.472 (0.134) | 0.589 (0.124) | 0.609 (0.112) | 0.870 (0.0903) | 0.514 (0.132) | 0.637 (0.116) | 0.356 (0.129) | 0.589 (0.222) | 0.369 (0.230) | 0.412 (0.175) | 0.442 (0.173) |
| ClaRNA | 0.808 (0.0931) | 0.449 (0.144) | 0.565 (0.128) | 0.589 (0.113) | 0.864 (0.0770) | 0.487 (0.139) | 0.612 (0.121) | 0.336 (0.0911) | 0.613 (0.232) | 0.343 (0.222) | 0.392 (0.149) | 0.430 (0.150) |
| DSSR | 0.769 (0.113) | 0.292 (0.0966) | 0.419 (0.111) | 0.465 (0.104) | 0.856 (0.0969) | 0.322 (0.111) | 0.459 (0.121) | 0.243 (0.109) | 0.462 (0.329) | 0.170 (0.175) | 0.234 (0.203) | 0.266 (0.210) |
| MC-Annotate | **0.883** (0.0743) | 0.335 (0.113) | 0.475 (0.123) | 0.531 (0.0992) | **0.917** (0.0640) | 0.365 (0.114) | 0.511 (0.120) | 0.318 (0.0701) | **0.709** (0.257) | 0.215 (0.114) | 0.321 (0.154) | 0.381 (0.160) |

**Table S13. Performance comparison between the geometric attention-based neural networks with and without parameterized geometric normalization for base-pair prediction.** Values are presented as mean (standard deviation). For each metric, the higher values between the two models shown in bold.

| Prediction mode | Dataset | Structure | With parameterized geometric normalization | Without parameterized geometric normalization |
|---|---|---|---|---|
| Backbone frame-based prediction | Benchmark dataset | Native | 0.890 (0.0933) | **0.898** (0.0841) |
| | | AlphaFold3 | **0.724** (0.258) | 0.717 (0.257) |
| | | DRfold2 | **0.725** (0.221) | 0.716 (0.222) |
| | Cryo-RM dataset | Native | 0.917 (0.0532) | **0.924** (0.0499) |
| | | EMRNA | 0.653 (0.181) | **0.654** (0.170) |
| C4'-trace-based prediction | Benchmark dataset | Native | **0.834** (0.124) | 0.737 (0.165) |
| | | AlphaFold3 | **0.710** (0.252) | 0.670 (0.233) |
| | | DRfold2 | **0.709** (0.220) | 0.665 (0.208) |
| | Cryo-RM dataset | Native | **0.887** (0.0635) | 0.863 (0.0821) |
| | | EMRNA | **0.745** (0.169) | 0.702 (0.152) |

**Table S14. Performance comparison between the geometric attention-based neural networks with and without parameterized geometric normalization for base-stacking prediction.** Values are presented as mean (standard deviation). For each metric, the higher values between the two models shown in bold.

| Prediction mode | Dataset | Structure | With parameterized geometric normalization | Without parameterized geometric normalization |
|---|---|---|---|---|
| Backbone frame-based prediction | Benchmark dataset | Native | 0.895 (0.0388) | **0.901** (0.0347) |
| | | AlphaFold3 | **0.786** (0.0942) | 0.784 (0.0948) |
| | | DRfold2 | 0.784 (0.0912) | **0.786** (0.0870) |
| | Cryo-RM dataset | Native | 0.909 (0.0475) | **0.910** (0.0454) |
| | | EMRNA | **0.589** (0.124) | 0.584 (0.125) |
| C4'-trace-based prediction | Benchmark dataset | Native | **0.806** (0.0600) | 0.791 (0.0668) |
| | | AlphaFold3 | **0.754** (0.0876) | 0.750 (0.0769) |
| | | DRfold2 | **0.748** (0.0896) | 0.739 (0.0866) |
| | Cryo-RM dataset | Native | **0.849** (0.0607) | 0.843 (0.0552) |
| | | EMRNA | **0.759** (0.102) | 0.727 (0.103) |

**Table S15. Comparison of refinement performance among the DRfold2 coarse-grained structure, the initial full-atom structure, refinement using Monte Carlo (MC) simulation only, refinement using L-BFGS energy minimization only, and refinement using a combination of MC simulation and L-BFGS energy minimization (RNArefine). The performance was evaluated using 17 types of metrics.**

| | RMSD | TM-score | Penalized TM-score | Clashscore | lDDT | Penalized lDDT | RMSD (Bond length) | RMSD (Bond angle) | MolProbity score | HB-score | MCQ | INF_all | INF_stack | INF_wc | INF_nwc | GDT_TS | Composite score |
|---|---|---|---|---|---|---|---|---|---|---|---|---|---|---|---|---|---|
| DRfold2 coarse-grained structure | **12.5** | 0.422 | - | - | - | - | - | - | - | - | - | - | - | - | - | 0.484 | - |
| Initial full-atomic structure | 12.5 | 0.421 | 0.127 | 116 | 0.634 | 0.0799 | 0.0470 | 8.28 | 3.64 | 0.506 | 21.4 | 0.755 | 0.766 | **0.816** | 0.437 | 0.484 | -3.31 |
| MC only | 12.5 | **0.422** | 0.408 | 26.8 | **0.636** | 0.608 | 0.0204 | 2.08 | 2.92 | 0.508 | 21.8 | 0.773 | 0.785 | 0.815 | **0.497** | **0.484** | 0.808 |
| L-BFGS only | 12.6 | 0.421 | 0.419 | 11.1 | 0.635 | 0.628 | 0.0195 | 1.71 | 2.40 | 0.532 | **20.8** | 0.773 | 0.790 | 0.815 | 0.476 | 0.483 | 1.19 |
| MC+L-BFGS (RNArefine) | 12.6 | 0.422 | **0.420** | **8.16** | 0.634 | **0.628** | **0.0143** | **1.67** | **2.26** | **0.533** | 21.1 | **0.775** | **0.790** | 0.815 | 0.486 | 0.483 | **1.31** |

**Table S16. Comparison of physical metrics between native structures and structures refined by RNArefine in the benchmark on our dataset.**

| | Clashscore | RMSD (Bond length) | RMSD (Bond angle) | MolProbity score |
|---|---|---|---|---|
| Native | 5.40 | 0.00514 | 0.903 | 2.27 |
| RNArefine | 8.16 | 0.0143 | 1.67 | 2.26 |

**Table S17. Comparison of RNArefine performance with and without predicted interaction information in the benchmark dataset.** RNArefine was evaluated with and without energy terms derived from predicted base-pair and base-stacking interactions.

| | RMSD | TM-score | Penalized TM-score | Clashscore | lDDT | Penalized lDDT | RMSD (Bond length) | RMSD (Bond angle) | MolProbity score | HB-score | MCQ | INF_all | INF_stack | INF_wc | INF_nwc | GDT_TS |
|---|---|---|---|---|---|---|---|---|---|---|---|---|---|---|---|---|
| RNArefine (no interaction) | **12.5** | 0.417 | 0.416 | **3.12** | 0.572 | 0.570 | 0.0146 | 1.73 | **1.90** | 0.202 | 25.4 | 0.659 | 0.710 | 0.608 | 0.227 | 0.481 |
| RNArefine | 12.6 | **0.422** | **0.420** | 8.16 | **0.634** | **0.628** | **0.0143** | **1.67** | 2.26 | **0.533** | **21.1** | **0.775** | **0.790** | **0.815** | **0.486** | **0.483** |

**Table S18. Comparison of the refinement performance of RNArefine with existing RNA refinement methods.** The performance was evaluated using 17 types of metrics.

| | RMSD | TM-score | Penalized TM-score | Clashscore | lDDT | Penalized lDDT | RMSD (Bond length) | RMSD (Bond angle) | MolProbity score | HB-score | MCQ | INF_all | INF_stack | INF_wc | INF_nwc | GDT_TS | Composite score |
|---|---|---|---|---|---|---|---|---|---|---|---|---|---|---|---|---|---|
| PDBfixer | **12.5** | **0.422** | 0.00101 | 196 | 0.535 | 0.00 | 0.0363 | 12.4 | 3.89 | 0.0556 | 28.9 | 0.447 | 0.494 | 0.352 | 0.0375 | 0.484 | -9.05E+13 |
| RNAfitme | 12.5 | 0.421 | 0.130 | 63.4 | 0.631 | 0.0815 | 0.0619 | 8.51 | 3.30 | 0.330 | 22.0 | 0.766 | 0.777 | 0.812 | 0.459 | **0.484** | -1.52 |
| Arena | 12.5 | 0.421 | 0.254 | 42.3 | **0.634** | 0.258 | 0.0422 | 5.89 | 3.21 | 0.508 | 23.6 | 0.766 | 0.781 | **0.817** | 0.424 | 0.483 | -0.226 |
| QRNAS | 12.7 | 0.411 | 0.373 | 46.4 | 0.621 | 0.539 | 0.0437 | 3.85 | 2.45 | 0.367 | 21.6 | 0.772 | 0.786 | 0.814 | 0.479 | 0.475 | 0.635 |
| MD | 12.9 | 0.390 | 0.382 | 8.37 | 0.561 | 0.541 | 0.0280 | 2.54 | **2.16** | 0.519 | 26.7 | 0.754 | 0.769 | 0.799 | 0.428 | 0.445 | -7.73 |
| RNArefine | 12.6 | 0.422 | **0.420** | **8.16** | **0.634** | **0.628** | **0.0143** | **1.67** | 2.26 | **0.533** | **21.1** | **0.775** | **0.790** | 0.815 | **0.486** | 0.483 | **2.57** |

**Table S19. Comparison of RNArefine with existing RNA refinement methods on structures predicted by EMRNA from cryo-EM density maps.** Performance was evaluated using 17 metrics.

| | RMSD | TM-score | Penalized TM-score | Clashscore | lDDT | Penalized lDDT | RMSD (Bond length) | RMSD (Bond angle) | MolProbity score | HB-score | MCQ | INF_all | INF_stack | INF_wc | INF_nwc | GDT_TS | Composite score |
|---|---|---|---|---|---|---|---|---|---|---|---|---|---|---|---|---|---|
| Initial structure | 2.67 | 0.705 | 0.0806 | 176 | 0.601 | 0.0178 | 0.0969 | 10.0 | 3.84 | 0.380 | 41.4 | 0.607 | 0.599 | 0.751 | 0.208 | 0.730 | -4.27 |
| QRNAS | 2.51 | 0.720 | 0.548 | 80.6 | 0.632 | 0.473 | 0.0745 | 5.57 | 2.87 | 0.284 | **39.4** | 0.679 | 0.684 | 0.775 | 0.241 | 0.749 | 0.240 |
| MD | 2.68 | 0.687 | 0.620 | 28.2 | 0.567 | 0.505 | 0.0483 | 3.26 | 2.66 | 0.385 | 45.2 | 0.653 | 0.659 | 0.762 | 0.196 | 0.700 | -0.805 |
| RNArefine (C4'-trace) | **2.47** | **0.731** | **0.722** | **5.53** | **0.648** | **0.634** | **0.0182** | 2.12 | **2.26** | **0.471** | 40.6 | **0.757** | **0.760** | **0.866** | **0.315** | **0.757** | **3.26** |
| RNArefine (Backbone) | 2.51 | 0.727 | 0.715 | 9.04 | 0.587 | 0.573 | 0.0251 | **2.09** | 2.37 | 0.407 | 42.8 | 0.661 | 0.659 | 0.783 | 0.250 | 0.753 | 1.58 |

**Table S20. Comparison of physical metrics between native structures and structures refined by RNArefine in the benchmark on Cryo-EM dataset.**

| | Clashscore | RMSD (Bond length) | RMSD (Bond angle) | MolProbity score |
|---|---|---|---|---|
| Native | 4.59 | 0.00477 | 0.762 | 2.28 |
| RNArefine (C4'-trace) | 5.53 | 0.0182 | 2.12 | 2.26 |
| RNArefine (Backbone) | 9.04 | 0.0251 | 2.09 | 2.37 |

**Table S21. Prediction performance of the top 30 participating groups and refined models on the 16 RNA monomer targets in CASP16.** Z-scores were computed by the official CASP16 RNA monomer evaluation protocol. Performance metrics for the original submitted models were retrieved from source data obtained from the official CASP16 website, whereas metrics for the refined structures (in bold fonts) were estimated in this study. In addition to the Z-score, TM-score, Penalized lDDT, and GDT-TS were averaged over all targets.

| Group | Z-Score | TM-score | Penalized lDDT | GDT-TS |
|---|---|---|---|---|
| **Refined_Vfold** | 10.6 | 0.449 | 0.654 | 0.430 |
| Vfold | 10.5 | 0.451 | 0.655 | 0.430 |
| **Refined_GuangzhouRNA-human** | 10.2 | 0.457 | 0.667 | 0.429 |
| **Refined_GuangzhouRNA-meta** | 9.67 | 0.403 | 0.638 | 0.376 |
| **Refined_Bhattacharya** | 9.44 | 0.403 | 0.654 | 0.386 |
| **Refined_KiharaLab** | 9.29 | 0.439 | 0.659 | 0.418 |
| **Refined_BRIQX** | 9.00 | 0.474 | 0.676 | 0.463 |
| **Refined_AF3-server** | 8.99 | 0.417 | 0.661 | 0.395 |
| **Refined_CSSB_experimental** | 8.98 | 0.435 | 0.640 | 0.405 |
| KiharaLab | 8.76 | 0.442 | 0.653 | 0.422 |
| **Refined_elofsson** | 8.67 | 0.422 | 0.663 | 0.405 |
| GuangzhouRNA-human | 8.52 | 0.460 | 0.636 | 0.431 |
| CSSB_experimental | 8.38 | 0.434 | 0.631 | 0.404 |
| BRIQX | 7.93 | 0.476 | 0.655 | 0.466 |
| GuangzhouRNA-meta | 7.51 | 0.403 | 0.600 | 0.377 |
| AF3-server | 7.47 | 0.418 | 0.638 | 0.395 |
| **Refined_Yang-Server** | 7.46 | 0.432 | 0.628 | 0.403 |
| **Refined_RNAFOLDX** | 7.35 | 0.379 | 0.619 | 0.359 |
| **Refined_LCBio** | 7.27 | 0.492 | 0.667 | 0.460 |
| LCBio | 7.23 | 0.496 | 0.660 | 0.461 |
| **Refined_Diff** | 6.93 | 0.336 | 0.522 | 0.318 |
| elofsson | 6.83 | 0.422 | 0.643 | 0.405 |
| Yang-Server | 6.65 | 0.431 | 0.618 | 0.402 |
| **Refined_GromihaLab** | 6.57 | 0.406 | 0.641 | 0.419 |
| **Refined_Zheng** | 6.25 | 0.400 | 0.621 | 0.386 |
| GromihaLab | 6.19 | 0.405 | 0.619 | 0.420 |
| **Refined_NKRNA-s** | 6.01 | 0.392 | 0.613 | 0.379 |

| | | | | |
|---|---|---|---|---|
| Diff | 5.80 | 0.326 | 0.507 | 0.306 |
| Bhattacharya | 5.76 | 0.405 | 0.437 | 0.385 |
| **Refined_Yang-Multimer** | 5.64 | 0.426 | 0.631 | 0.432 |
| NKRNA-s | 5.54 | 0.392 | 0.596 | 0.380 |
| RNAFOLDX | 5.53 | 0.378 | 0.591 | 0.357 |
| **Refined_dNAfold** | 5.50 | 0.407 | 0.605 | 0.408 |
| CoDock | 5.17 | 0.488 | 0.690 | 0.481 |
| **Refined_CoDock** | 5.17 | 0.485 | 0.684 | 0.477 |
| Yang-Multimer | 5.15 | 0.426 | 0.619 | 0.434 |
| **Refined_MIEnsembles-Server** | 5.06 | 0.385 | 0.605 | 0.367 |
| Zheng | 4.93 | 0.400 | 0.606 | 0.386 |
| **Refined_RNApolis** | 4.75 | 0.393 | 0.617 | 0.391 |
| MIEnsembles-Server | 4.63 | 0.386 | 0.587 | 0.368 |
| RNApolis | 4.44 | 0.394 | 0.610 | 0.392 |
| **Refined_falcon2** | 4.42 | 0.367 | 0.620 | 0.356 |
| **Refined_B-LAB** | 4.35 | 0.320 | 0.518 | 0.309 |
| **Refined_OpenComplex** | 4.26 | 0.322 | 0.546 | 0.307 |
| **Refined_OpenComplex_Server** | 4.20 | 0.288 | 0.521 | 0.266 |
| **Refined_406** | 3.88 | 0.316 | 0.527 | 0.289 |
| **Refined_GeneSilico** | 3.61 | 0.382 | 0.580 | 0.377 |
| 406 | 3.58 | 0.313 | 0.509 | 0.283 |
| B-LAB | 3.54 | 0.320 | 0.494 | 0.308 |
| OpenComplex | 3.17 | 0.319 | 0.516 | 0.303 |
| RNA_Dojo | 3.16 | 0.364 | 0.595 | 0.375 |
| falcon2 | 3.15 | 0.366 | 0.553 | 0.356 |
| GeneSilico | 3.15 | 0.381 | 0.570 | 0.377 |
| **Refined_dfr** | 2.97 | 0.383 | 0.648 | 0.418 |
| **Refined_405** | 2.96 | 0.287 | 0.519 | 0.263 |
| 405 | 2.86 | 0.284 | 0.501 | 0.258 |
| **Refined_RNA_Dojo** | 2.84 | 0.361 | 0.598 | 0.373 |
| OpenComplex_Server | 2.83 | 0.288 | 0.473 | 0.265 |
| dNAfold | 2.44 | 0.408 | 0.361 | 0.406 |
| dfr | 1.52 | 0.382 | 0.518 | 0.420 |

**Table S22. Z-scores of the top 30 participating groups and refined models on the 16 RNA monomer targets in CASP16.** The table reports the sum of Z-scores for lDDT, TM-score, and GDT-TS, as well as the total sum of Z-scores and the sum of positive Z-scores **(score in CASP16)**. For each metric, Z-scores were summed across all targets for each group. Original and refined models are compared for each score, and the higher values shown in bold.

| Group | Sum of Z-Score (penalized lDDT) | | Sum of Z-Score (TM-score) | | Sum of Z-Score (GDT-TS) | | Sum of Z-Score | | Sum of positive Z-Score | |
|---|---|---|---|---|---|---|---|---|---|---|
| | Decoy | Refined | Decoy | Refined | Decoy | Refined | Decoy | Refined | Decoy | Refined |
| Vfold | 12.6 | **12.9** | **10.7** | 10.4 | 12.1 | 12.1 | 9.98 | **10.1** | 10.5 | **10.6** |
| KiharaLab | 11.6 | **13.2** | **8.53** | 8.10 | **9.70** | 8.90 | 8.33 | **8.80** | 8.76 | **9.29** |
| GuangzhouRNA-human | 8.97 | **14.2** | **9.33** | 9.05 | **12.2** | 12.1 | 8.45 | **10.1** | 8.52 | **10.2** |
| CSSB_experimental | 10.1 | **11.5** | 9.52 | **10.0** | 8.05 | **8.51** | 7.69 | **8.51** | 8.38 | **8.98** |
| BRIQX | 9.24 | **12.3** | **7.35** | 7.29 | **8.38** | 8.17 | 6.85 | **7.88** | 7.93 | **9.00** |
| GuangzhouRNA-meta | 5.80 | **12.3** | 7.72 | **7.85** | **6.59** | 6.12 | 6.82 | **9.13** | 7.51 | **9.67** |
| AF3-server | 9.78 | **14.6** | **6.81** | 6.68 | **6.62** | 5.99 | 7.07 | **8.75** | 7.47 | **8.99** |
| LCBio | 7.83 | **9.51** | **8.49** | 8.08 | 7.46 | **7.53** | 6.57 | **7.20** | 7.23 | **7.27** |
| elofsson | 10.1 | **14.5** | **5.23** | 5.11 | **8.39** | 7.90 | 6.75 | **8.43** | 6.83 | **8.67** |
| Yang-Server | 7.55 | **9.00** | 7.32 | **7.78** | 6.40 | **7.14** | 6.50 | **7.34** | 6.65 | **7.46** |
| GromihaLab | 4.82 | **7.28** | 3.18 | **3.20** | **5.29** | 5.09 | 4.22 | **4.91** | 6.19 | **6.57** |
| Diff | -5.49 | **-2.51** | -9.57 | **-7.60** | -9.86 | **-7.62** | -7.76 | **-5.42** | 5.80 | **6.93** |
| Bhattacharya | -13.5 | **11.4** | **7.50** | 7.20 | 5.78 | **6.13** | 0.644 | **8.79** | 5.76 | **9.44** |
| NKRNA-s | 3.04 | **7.07** | 0.545 | **0.848** | **3.28** | 3.13 | 2.74 | **4.43** | 5.54 | **6.01** |
| RNAFOLDX | 3.89 | **8.81** | 2.04 | **2.45** | **4.40** | 4.37 | 3.85 | **5.71** | 5.53 | **7.35** |
| CoDock | **7.67** | 7.53 | **5.54** | 5.29 | **5.80** | 5.37 | **4.96** | 4.80 | 5.17 | 5.17 |
| Yang-Multimer | 4.30 | **5.89** | 5.77 | **6.11** | **4.32** | 4.01 | 4.11 | **4.65** | 5.15 | **5.64** |
| Zheng | 4.94 | **8.57** | 1.80 | **2.10** | 4.26 | **4.90** | 2.77 | **4.60** | 4.93 | **6.25** |
| MIEnsembles-Server | 2.12 | **6.46** | -0.0189 | **0.0441** | **1.32** | 1.05 | 2.02 | **3.66** | 4.63 | **5.06** |
| RNApolis | 5.04 | **5.98** | 0.590 | **0.634** | **-2.31** | -2.33 | 1.27 | **1.68** | 4.44 | **4.75** |
| 406 | -2.18 | **0.545** | -3.61 | **-3.21** | -4.77 | **-4.34** | -3.39 | **-2.05** | 3.58 | **3.88** |
| B-LAB | -7.62 | **-3.84** | **-10.7** | -11.0 | -8.07 | **-7.95** | -5.49 | **-4.11** | 3.54 | **4.35** |
| OpenComplex | -5.95 | **-1.18** | -10.3 | **-9.23** | -11.8 | **-10.8** | -7.24 | **-4.93** | 3.17 | **4.26** |
| RNA_Dojo | 0.646 | **1.17** | **-7.26** | -7.50 | -8.33 | **-8.25** | -3.68 | **-3.44** | **3.16** | 2.84 |
| falcon2 | -0.839 | **7.02** | -2.10 | **-2.00** | **-0.739** | -0.810 | -0.501 | **2.53** | 3.15 | **4.42** |
| GeneSilico | 1.48 | **2.93** | -1.82 | **-1.5** | **-5.67** | -5.80 | -3.47 | **-2.75** | 3.15 | **3.61** |
| 405 | -2.07 | **0.281** | -3.32 | **-3.13** | **-3.78** | -3.84 | -2.96 | **-1.98** | 2.86 | **2.96** |
| OpenComplex_Server | -10.3 | **-4.53** | -14.9 | **-14.5** | **-16.9** | -17.1 | -11.4 | **-9.17** | 2.83 | **4.20** |
| dNAfold | -27.6 | **2.33** | **0.435** | 0.126 | -2.35 | **-1.83** | -9.83 | **-0.237** | 2.44 | **5.50** |
| dfr | -6.76 | **5.21** | -2.01 | **-1.84** | **-0.578** | -0.853 | -3.16 | **1.48** | 1.52 | **2.97** |

**Table S23. Number of base-pair and non-base-pair interactions for each nucleotide pair in our training dataset.**

| Nucleotide pair | Base-pair interactions | Non-base-pair interactions |
|---|---|---|
| A-A | 2439 | 138030 |
| A-U | 20359 | 228464 |
| A-G | 7969 | 332253 |
| A-C | 2034 | 263114 |
| U-U | 1172 | 103786 |
| U-G | 7923 | 272523 |
| U-C | 935 | 214128 |
| G-G | 1702 | 209921 |
| G-C | 42137 | 334854 |
| C-C | 554 | 129161 |

**Table S24. Number of base-stacking and non–base-stacking interactions for each nucleotide pair in our training dataset.**

| Nucleotide pair | Adjacent | | Non-adjacent | |
|---|---|---|---|---|
| | Stacking | Non-Stacking | Stacking | Non-Stacking |
| A-A | 15051 | 9465 | 6318 | 64393 |
| A-U | 9288 | 5934 | 3888 | 123877 |
| A-G | 14684 | 8118 | 12443 | 159633 |
| A-C | 11065 | 4941 | 4075 | 127582 |
| U-A | 6001 | 8873 | - | - |
| U-U | 9203 | 8936 | 466 | 52610 |
| U-G | 10024 | 9704 | 4896 | 137476 |
| U-C | 9482 | 6592 | 977 | 106526 |
| G-A | 12113 | 10860 | - | - |
| G-U | 13590 | 5825 | - | - |
| G-G | 24744 | 5097 | 10374 | 94730 |
| G-C | 17947 | 4137 | 10034 | 168324 |
| C-A | 9392 | 7034 | - | - |
| C-U | 11755 | 4404 | - | - |
| C-G | 12694 | 8212 | - | - |
| C-C | 16075 | 6011 | 370 | 61270 |

### Supplementary Figures

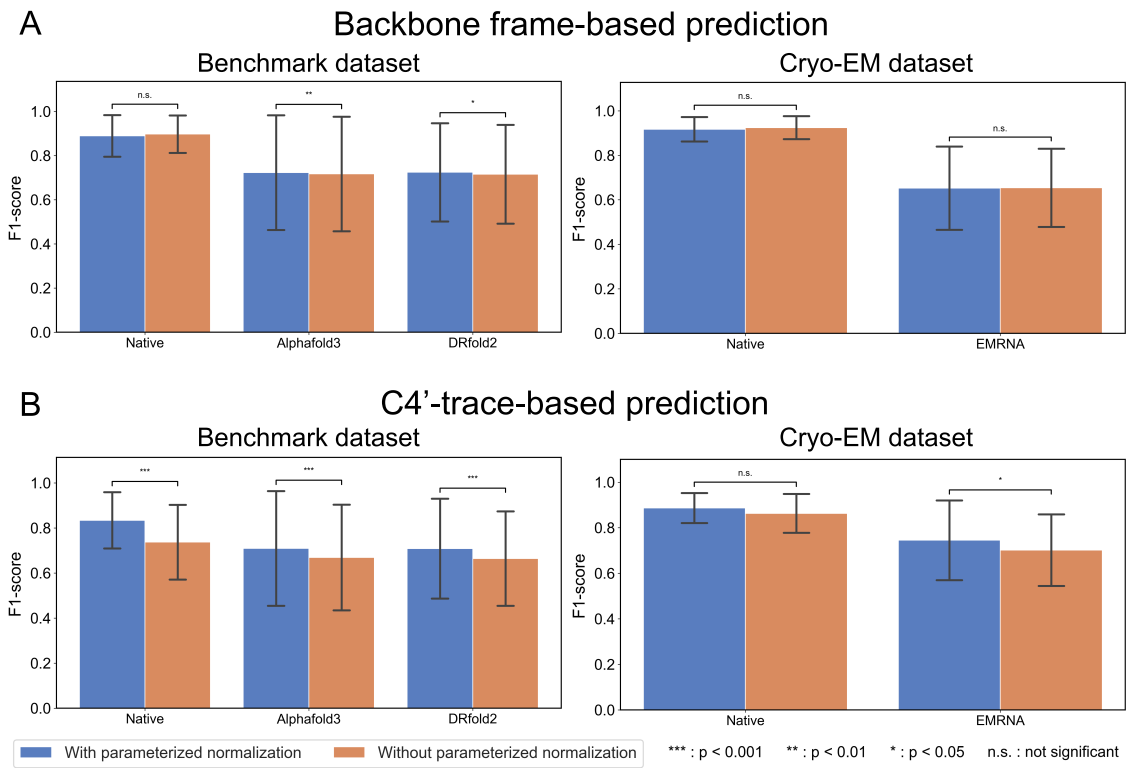

**Fig. S1 Performance comparison between the geometric attention-based neural networks with and without parameterized geometric normalization for base-pair prediction.** (A) Backbone-frame-based prediction in our benchmark dataset (left) and Cyro-EM dataset (right). (B) C4'-trace-based prediction in our benchmark dataset (left) and Cyro-EM dataset (right). To evaluate the effect of parameterized geometric normalization, prediction accuracy was compared between the two models using sample-wise one-sided t-tests.

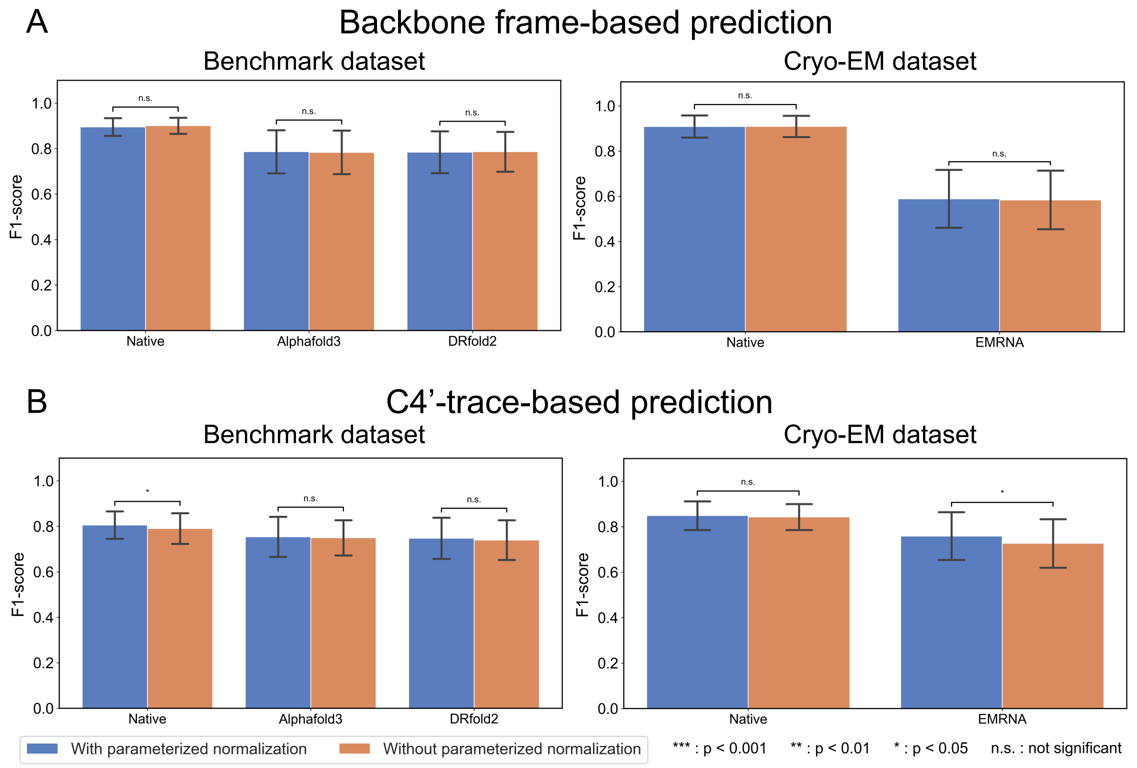

**Fig. S2 Performance comparison between the geometric attention-based neural networks with and without parameterized geometric normalization for base-stacking prediction.** (A) Backbone-frame-based prediction in our benchmark dataset (left) and Cyro-EM dataset (right). (B) C4'-trace-based prediction in our benchmark dataset (left) and Cyro-EM dataset (right). To evaluate the effect of parameterized geometric normalization, prediction accuracy was compared between the two models using sample-wise one-sided t-tests.

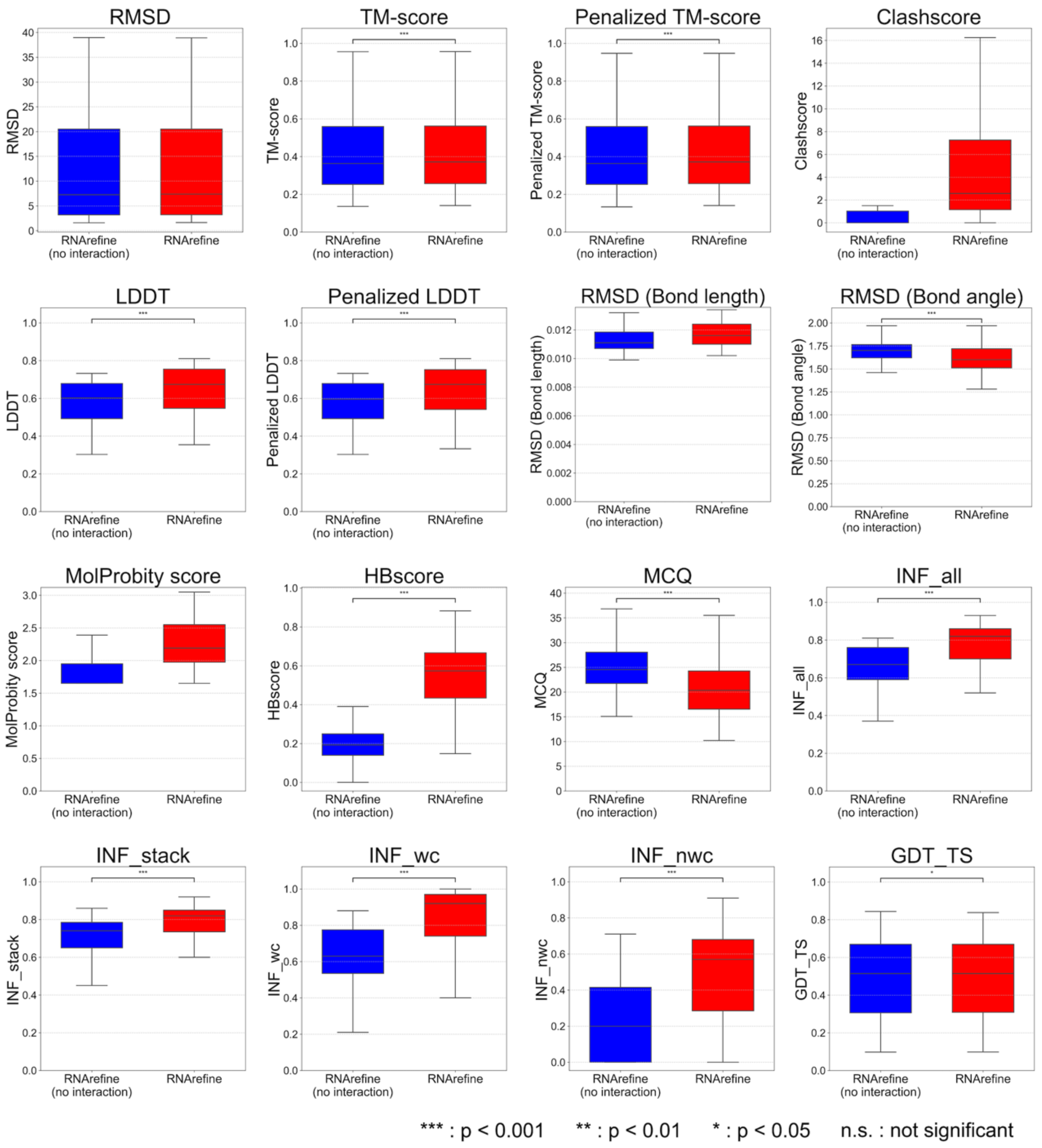

**Fig. S3 Comparison of RNArefine performance with and without predicted interaction information in our benchmark dataset.** RNArefine was evaluated with (red) and without (blue) energy terms derived from predicted base-pair and base-stacking interactions. Statistical significance was assessed using a one-sided sample-wise t-test to evaluate whether the inclusion of predicted interactions improves performance.

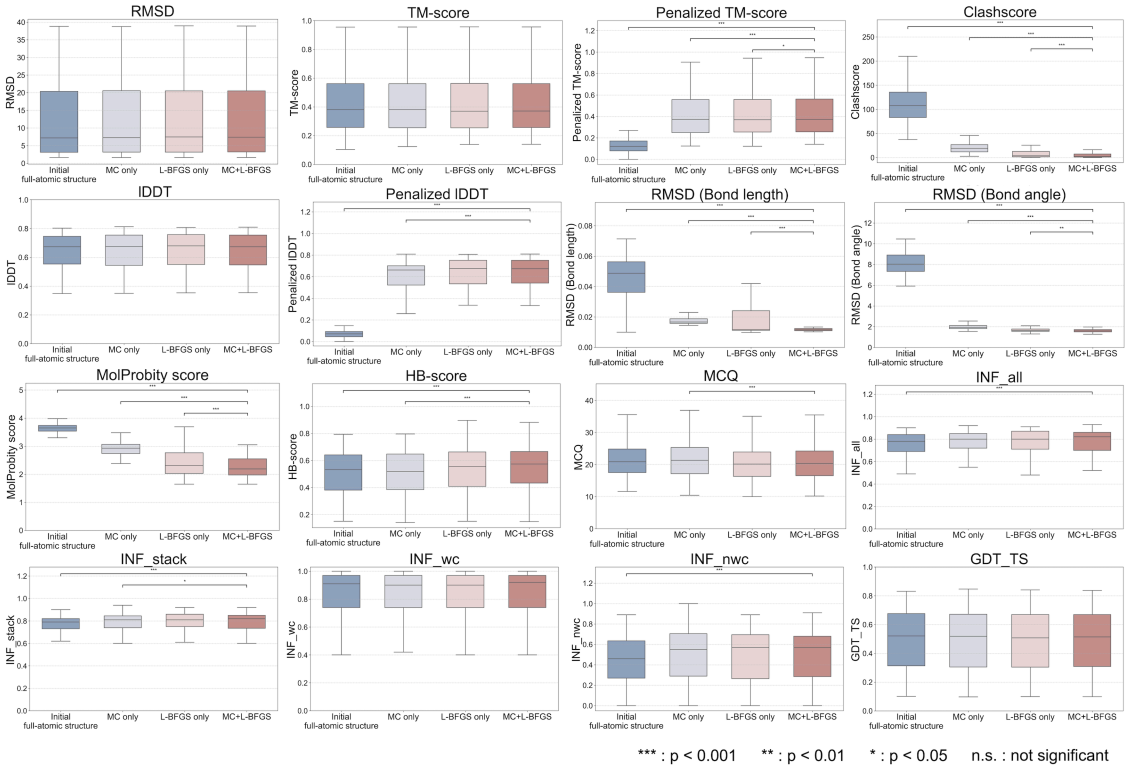

**Fig. S4 Refinement performance comparison among the initial full-atom structures, the fully refined structures (MC+L-BFGS), and the structures refined by each step individually (MC only and L-BFGS only).** For each refinement condition, sixteen evaluation metrics were calculated. Statistical significance was assessed for each metric using sample-wise one-sided t-tests, comparing the fully refined structures with each of the other conditions.

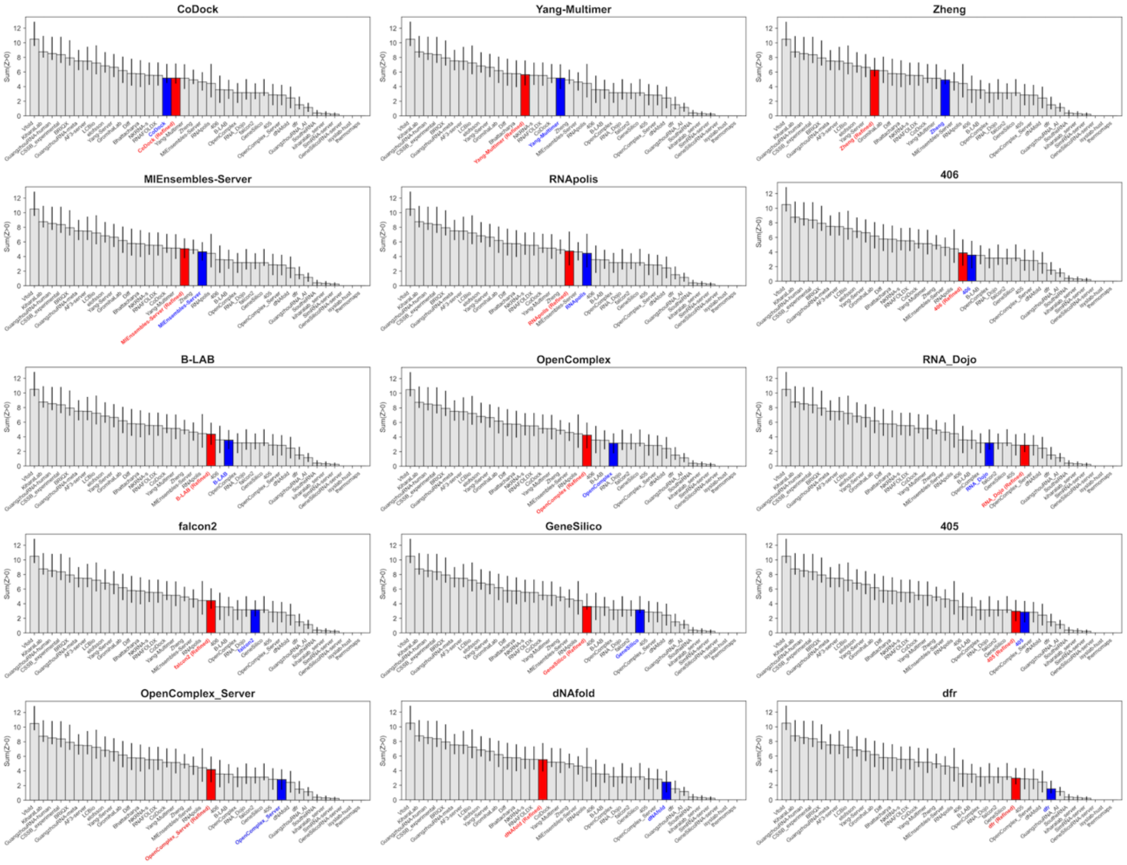

**Fig. S5 Improvement in RNA structure prediction performance by refinement in CASP16 (participant groups ranked 16–30 across 16 targets).** Changes in CASP rankings before and after refinement are shown for the participating groups ranked 16–30. Each panel corresponds to a participant group and displays the summed Z-score ranking across groups based on the originally submitted models (red) and the estimated ranking after refinement (blue). Panels include the top 30 groups, with the lower panels representing groups ranked 16–30. Z-scores and rankings were computed following the CASP assessment procedure, with confidence intervals estimated by bootstrap resampling using code adapted from the publicly available CASP16 nucleic acid assessment scripts (Kretsch, R. C. et al. Assessment of Nucleic Acid Structure Prediction in CASP16. bioRxiv 2025.05.06.652459, 2025)
